# Modular Derivation and Unbiased Single-cell Analysis of Regional Human Hindbrain And Spinal Neurons Enables Discovery of Nuanced Transcriptomic Patterns along Developmental Axes

**DOI:** 10.1101/2021.10.14.464440

**Authors:** Nisha R. Iyer, Junha Shin, Stephanie Cuskey, Yucheng Tian, Noah R. Nicol, Tessa E. Doersch, Sunnie Grace McCalla, Sushmita Roy, Randolph S. Ashton

**Affiliations:** Wisconsin Institute for Discovery, University of Wisconsin-Madison; Department of Biomedical Engineering, University of Wisconsin-Madison; Department of Biostatistics and Medical Informatics, University of Wisconsin-Madison

## Abstract

Our inability to derive the vast neuronal diversity of the posterior central nervous system (pCNS) using human pluripotent stem cells (hPSCs) poses a major impediment to understanding human neurodevelopment and disease in the hindbrain and spinal cord. Here we establish a modular differentiation paradigm that recapitulates patterning along both the rostrocaudal (R/C) and dorsoventral (D/V) axes of the pCNS, enabling derivation of any neuronal phenotype with discrete regional specificity. First, neuromesodermal progenitors (NMPs) with discrete Hox profiles are efficiently converted to pCNS progenitors (pCNSPs). Then by tuning D/V signaling, pCNSPs are directed to ventral Shh-dependent MNs (MNs) and locomotor interneurons (INs) or dorsal TGF-β-dependent proprioceptive INs and TGF-β-independent sensory INs. We applied D/V protocols to NMPs spanning the R/C axis for expansive single-cell RNA-sequencing (scRNAseq) analysis. By implementing a novel computational pipeline comprising sparse non-negative matrix factorization, consensus clustering, and combinatorial gene expression pattern identification, we detect hundreds of transcriptional markers within region-specific neuronal phenotypes, enabling discovery of gene expression patterns along the developmental axes. These findings highlight the potential of these resources to advance a mechanistic understanding of pCNS development, expand the potential and accuracy of *in vitro* models, and inform novel regenerative therapeutic strategies.

## Introduction

Nervous system diversity arises in response to a complex choreography of spatiotemporally restricted cues along the elongating embryo’s rostrocaudal (R/C) and dorsoventral (D/V) axes. These coordinated patterning events encode neural progenitors and post-mitotic neurons with unique transcriptional signatures that define a myriad of subtypes, which in turn orchestrate the precise neural circuits that shape human behavior (Lu et al., 2015; Philippidou and Dasen, 2013; Tanabe and Jessell, 1996). While hPSC-based approaches can, in theory, provide access to all these populations, differentiation strategies have intensely focused on recapitulating spinal D/V patterning with less attention to the generation of subtypes along the R/C axis. Even so, direct differentiation protocols have been achieved for relatively few cardinal neurons, which default to hindbrain or cervical identity (Amoroso et al., 2013; Butts et al., 2019; Du et al., 2015; Duval et al., 2019; Gupta et al., 2018; Maury et al., 2015). Motor neuron (MN) optimization has predominated, with robust differentiation schemas allowing high yields (Amoroso et al., 2013; Du et al., 2015; Maury et al., 2015) and some control over columnar and R/C identity (Ho et al., 2021; Maury et al., 2015; Mouilleau et al., 2021), but these protocols are not designed to adapt to other phenotypes. There has been some success in recreating R/C (Libby et al., 2021; Moris et al., 2020) and D/V (Andersen et al., 2020; Ogura et al., 2018; Zheng et al., 2019) signaling centers in human organoid models. However, the variability in efficiency, cell type distribution, and maturity of terminal populations, as well as the difficulty of cell recovery from organoid tissues, limits the scalability of these platforms for clinical applications.

We sought to develop a robust, modular differentiation methodology in monolayer culture to derive any posterior central nervous system (pCNS) phenotype by recapitulating the sequence of patterning events during development. Morphogenesis of the posterior neural tube, which forms the hindbrain and spinal cord (i.e. pCNS), begins near the primitive streak with a bipotent population of axial stem cells called neuromesodermal progenitors (NMPs) (Henrique et al., 2015; Wymeersch et al., 2016). As they proliferate, NMPs fuel R/C extension of the embryo, and their paraxial mesoderm or neuroectoderm progeny acquire a region-specific identity via combinatorial Hox gene expression (Deschamps and Duboule, 2017). The human genome contains 39 Hox genes subdivided into 13 paralogous groups (*HOX1-13*) arranged in four genomic clusters (*HOXA-D*). Maintenance of NMP bipotentiality—and thus progressive, colinear Hox gene activation— is governed by the balance between Wnt/β-catenin, fibroblast growth factor (FGF), and retinoic acid (RA) signaling pathways (Deschamps and Duboule, 2017; Diez Del Corral and Morales, 2017; Diez del Corral et al., 2003; Henrique et al., 2015; Liu, 2006; Liu et al., 2001; Neijts et al., 2016). A shift towards RA signaling prompts an exit from the bipotent NMP state to the neural fate and terminates Hox gene progression, resulting in neuroepithelial progeny with a precisely restricted R/C position via their HOX ‘code’ (Diez del Corral et al., 2003; Molotkova et al., 2005). Concurrent with folding of the neural plate to form the neural tube, D/V patterning is initiated by secretion of morphogens dorsally from the roof plate (bone morphogenetic proteins (BMPs) and Wnts) and ventrally from the floor plate (sonic hedgehog (Shh))(Wilson and Maden, 2005). These signals trigger concentration- and time-dependent expression of cross-repressive transcription factors that establish 11 discrete progenitor domains, broadly conferring somatosensory or locomotor phenotypes (Lai et al., 2016; Lu et al., 2015; Tanabe and Jessell, 1996). Though primarily considered drivers of R/C patterning, Hox genes remain dynamic through D/V specification and become restricted to discrete dorsal or ventral domains that correlate with the formation of distinct neuronal subtypes (Dasen and Jessell, 2009; Hayashi et al., 2018; Nolte and Krumlauf, 2013; Osseward et al., 2021; Philippidou and Dasen, 2013; Sweeney et al., 2018).

Previously, we showed that hPSCs could be efficiently converted to NMPs with discrete HOX profiles along the R/C axis by temporal modulation of Wnt, FGF, and RA signaling (Lippmann et al., 2015). Here, we expand on that work to demonstrate an optimized transition from the NMP to the pCNS progenitor state, enabling concentration and time dependent D/V patterning and rapid conversion to neurons with discrete regional phenotypes. We generated a single cell RNA-sequencing (scRNAseq) dataset comprising 59,502 cells that profile multiple points along the R/C and D/V axes, providing an expansive map of transcriptional programs that regulate neuronal specification. The novelty of our dataset also posed analytical challenges to neuronal characterization, as the reliance on known transcriptional markers determined from rodent development potentially excludes human-specific cell types. We established an unbiased cell population identification and characterization pipeline that identifies coarse-resolution primary clusters and fine-resolution subclusters corresponding to cell subtypes. Finally, we developed a strategy to characterize regionally or phenotypically comparable populations by identifying genes that exhibit combinatorial patterns of expression across cell types. Our computational analyses revealed differences in marker expression between our hPSC-derived neurons and embryonic mouse and human spinal neurons, novel expression patterns in cardinal neurons corresponding to different R/C positions, and evidence that perturbations in progenitor patterning persistently alter post-mitotic gene expression patterns. We anticipate that our modular differentiation paradigm and associated computational tools will be a valuable resource for biomanufacturing discrete, region-specific, pCNS populations, which will enable precise modeling of human development and disease as well as homologous cell grafts for regenerative medicine applications (Kadoya et al., 2016; Kumamaru et al., 2018).

## Results

### Smad Inhibition Optimizes Conversion of NMPs to Naive pCNS Progenitors

We first evaluated whether applying a single ventral patterning schema to hPSC-derived NMPs from diverse R/C regions would enable consistent derivation of ventral neuronal phenotypes. Using our HOX protocol, we derived six different NMP cultures from hESCs corresponding to 24hrs (H24), 48hrs (H48), 72hrs (H72), 120hrs (H120), 168hrs (H168), and 216hrs (H216) patterning periods in FGF8, CHIR, and/or GDF11 and dorsomorphin (**Fig. S1 A-B**)(Lippmann et al., 2015). NMP cultures were exposed to RA and small molecule Shh agonists Smoothened agonist (SAG) and Purmorphamine (Pur) prior to the addition of DAPT, which induces rapid neuronal conversion. Samples were cryopreserved, thawed, and cultured overnight prior to immunocytochemistry and scRNAseq analysis (**Fig. S1 B-D**).

In agreement with our prior publication, cultures expressed increasingly caudal HOX paralogs that could be correlated to cervical (*HOX1-8*; H24-vN, H48-vN, and H72-vN), thoracic (*HOX1-9*; H120-vN), lumbar (*HOX1-11*; H168-vN), and lumbosacral (*HOX1-13*; H216-vN) spinal regions (**Fig. S1E, Fig. S4A**)(Lippmann et al., 2015). We attributed the absence of hindbrain identities (expressing only *HOX1-4*) and similarity between H24, H48, and H72 Hox profiles to prolonged RA exposure during the neuronal differentiation stage, since RA alone is capable of caudalizing cells to a cervical fate (Mazzoni et al., 2013). Notably, analysis at single cell resolution revealed intra-sample uniformity in *HOX* expression (**Fig. S4A**). This illustrates our HOX protocol’s ability to discretize the pCNS R/C axis, in contrast to the broad or heterogenous *HOX* profiles observed in other direct differentiation protocols and organoid models (Duval et al., 2019; Gouti et al., 2014; Kumamaru et al., 2018; Libby et al., 2021; Ogura et al., 2018).

Though our aim was to produce cultures with high *SNAP25*^+^ neuronal content, cell type heterogeneity within and across samples was apparent by staining (**Fig. S1C**) and sample (**Fig. S1D**), cluster (**Fig. S1F**), and gene expression (**Fig. S1G, H**) distributions on tSNE visualizations of single cell transcriptomic data. Neural progenitor (*SOX2*^+^) and neuron (*SNAP25*^+^) composition varied between 10-80% (**Fig. S1I, J**). Thus, while samples could be patterned to discrete regions on the R/C axis, direct application of ventral morphogens caused inconsistent neuronal differentiation across different NMP populations.

We hypothesized that consistent neuronal differentiation from NMPs first requires efficient induction to a SOX2^+^/PAX6^+^ pCNS progenitors (pCNSPs), akin to the formation of neural plate epithelium from tail bud progenitors during gastrulation (Henrique et al., 2015). This process is regulated by RA and Noggin (a BMP antagonist) secreted by the somites and notochord (Diez del Corral et al., 2003; McMahon et al., 1998; Molotkova et al., 2005; Wilson et al., 2009). We derived SOX2^+^/T^+^ H120 NMPs (**Fig. S2 A**), then exposed them to RA and/or small molecule Smad inhibitors (SB+LDN) for up to 3 days (H120-pCNSPs, **Fig. S2B**). Both RA and SB+LDN were required to generate SOX2^+^/PAX6^+^ H120-pCNSPs efficiently (**Fig. S2C-Z**). In the absence of one or both factors, we observed persistent PAX3^+^ and PAX7^+^ cells that could become mesodermal (PAX3^+^/PAX7^+^), myogenic (PAX3^+^) (**Fig. S2C-T**) or neural crest (SOX10^+^) progeny (**Fig. S2GG-LL**). Both factors were also required to prevent inadvertent dorsal (PAX6^+^/PAX3^+^/PAX7^+^; AP2a^+^) (**Fig. S2O-T, S2GG-LL**), intermediate (PAX6^+^/PAX3^+^) (**Fig. S2O-T**) or ventral (NKX6.1^+^) (**Fig. S2AA-FF**) patterning. Thus, RA and SB+LDN cooperate to repress PAX3 and PAX7, which allows for the conversion of NMPs to unbiased, naive pCNSPs for subsequent D/V patterning.

### Concentration-Dependent Differentiation of pCNSPs Along D/V Axis

To simplify derivation of diverse pCNSPs with precise R/C positioning, we wanted to use the same ventralizing or dorsalizing differentiation schema for all cultures. Ventral interneurons (INs) and MNs arise in response to graded Shh signaling in the developing neural tube (Tanabe and Jessell, 1996; Wilson and Maden, 2005) (**Fig. 1A**). Thus, we first sought to determine whether hPSC-derived pCNSPs could be efficiently patterned to ventral identities in a concentration-dependent manner. We patterned H120-pCNSPs for 4 days in either 100nM or 1µM RA containing SB+LDN and varying concentrations of SAG and Pur to generate ventral progenitor cultures (**Fig. 1B**). Sustained exposure to SB+LDN suppressed PAX3 and PAX7 expression (**Fig. 1C-F**) while Shh signaling caused concentration-dependent increases in ventral progenitor markers (**Fig. 1G-M**). Notably, reducing the concentration of RA during ventral patterning improved the potency of Shh signaling, resulting in significant increases in *NKX6.1, OLIG2, and NKX2.2* expression in optimal culture conditions (**Fig. 1K-M**). Exposing ventral progenitor cultures to DAPT for 5 days induced rapid neuronal differentiation (**Fig. 1B**) and appropriately stratified post-mitotic INs and MNs (**Fig. 1A, N-Y**).

**Fig. 1.**
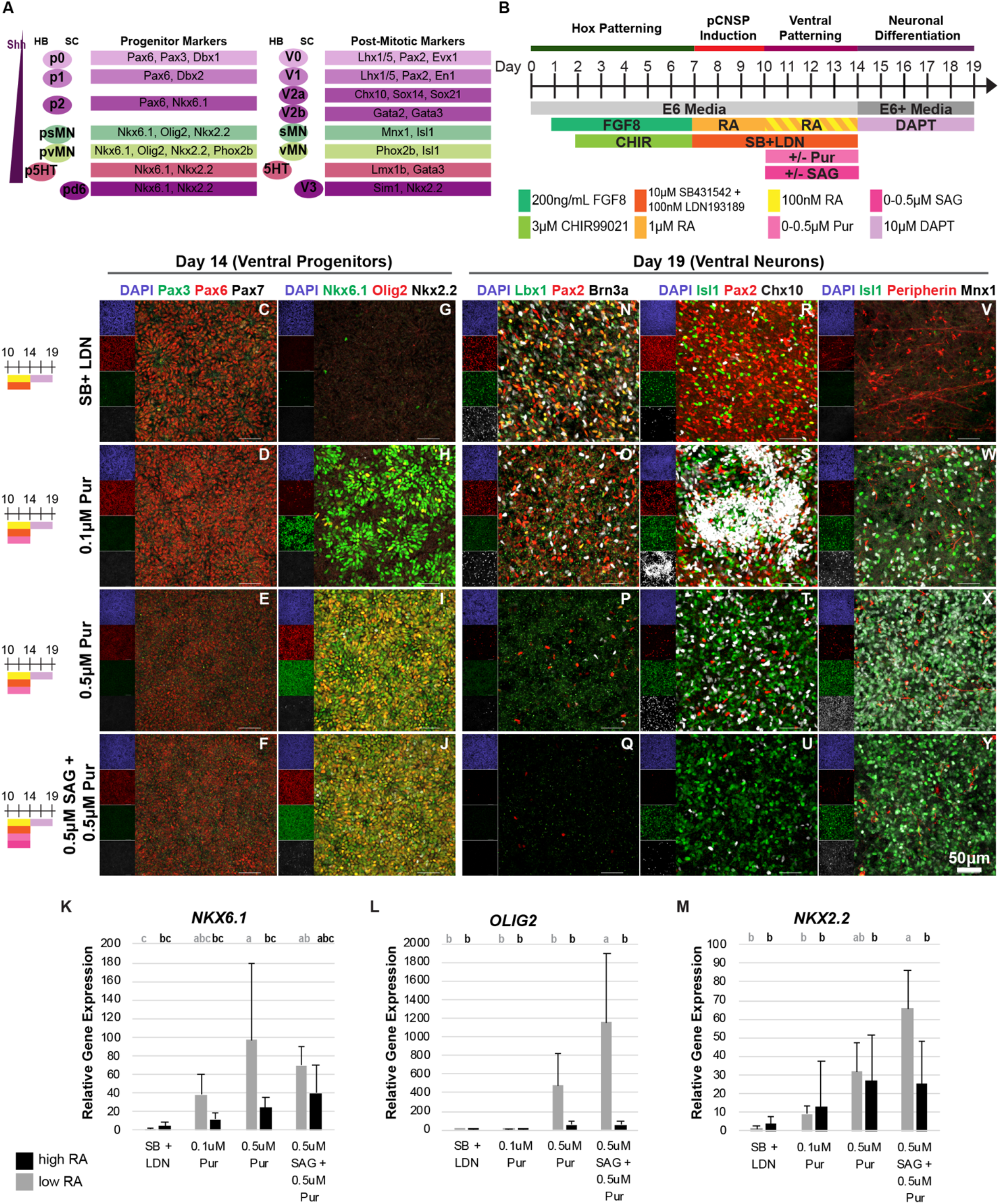
Concentration Dependent Shh Patterning of Ventral Spinal Neurons. (**A**) Ventral pCNS populations, with characteristic progenitor and post-mitotic transcription factor markers for the hindbrain (HB) and spinal cord (SC). (**B**) Timeline of ventral differentiation from H120-NMPs. (**C-J**) Immunostaining and (**K-M**) qRT-PCR in day 14 progenitor cultures. Error bars represent standard deviation (n=6 biological replicates per condition). Data shown as relative gene expression compared to 100nM RA SB+LDN condition. Statistics were calculated by one-way ANOVA with Tukey-Kramer post-hoc (samples with same letter assignment show no significant difference, p < 0.05). (**N-Y**) Immunostaining in day 19 post-mitotic neurons. All scale bars = 50μm. *See also Fig. S1 and S2*.

Efficient dorsal patterning of pCNS neurons *in vitro* has historically been difficult because of the ubiquitous roles of BMPs and Wnts elsewhere in the developing embryo. There has also been debate whether BMPs perform as morphogens (Duval et al., 2019; Tanabe and Jessell, 1996; Wilson and Maden, 2005) or act deterministically (Andrews et al., 2017; Gupta et al., 2018; Le Dreau et al., 2012; Lee et al., 1998), complicating efforts towards a streamlined differentiation strategy. To investigate this question, we cultured H120-pCNSPs for 4 days in either 100nM or 1µM RA containing Cyclopamine (Cyc)–a Shh antagonist–and varying concentrations and exposure durations of BMP4 to generate dorsal progenitor cultures (**Fig. 2B**). Termination of SB+LDN during dorsal patterning released suppression of PAX3 and PAX7 activity (**Fig. 2C-F**), which were elevated in response to increased BMP signaling **(Fig. 2L,M**). Importantly, this occurred without significant changes in *PAX6* expression, indicating the cells’ maintenance of a CNS identity (**Fig. 2K**). QRT-PCR-assessed gene expression patterns also indicated a shift from intermediate to dorsal fates with BMP4 exposure (**Fig. 2N,O**). While we observed OLIG3^+^ pd1/pd2/pd3 progenitors and the formation of some AP2α^+^ roof plate cells in conditions with the highest BMP4 exposure, no SOX10^+^ neural crest progeny was present (**Fig. 2J**). Treatment with DAPT induced rapid neuronal differentiation (**Fig. 2B**) and appropriately stratified post-mitotic dorsal INs in direct correlation to BMP4 exposure concentration/duration (**Fig. 2A**, **2P-AA**). This indicates that BMP4 behaves as a morphogen in agreement with other recent findings (Duval et al., 2019). Furthermore, because BMP7 has been shown to be required for neurogenesis of dI1/dI3/dI5 INs (Le Dreau et al., 2012), we wanted to determine whether adding BMP7 during the neuronal differentiation phase could push progenitors towards more dorsal post-mitotic fates (**Fig. 3A**). With BMP7 treatment, we observed a shift from dI4/dI5/dI6 to dI2/dI3 INs (**Fig. 3B,D,F**) and from dI2/dI3 to dI1/dI2 INs neurons (**Fig. 3C,E,G**) in progenitors pulsed or maintained in 20ng/mL BMP4 over the dorsal patterning period, respectively. Collectively, the results demonstrate our differentiation schema generates the full spectrum of D/V cell types from a single R/C position (H120), with the ability to obtain desired subtypes by optimizing morphogen exposure within discrete timeframes (**Fig. 3H**).

**Fig. 2.**
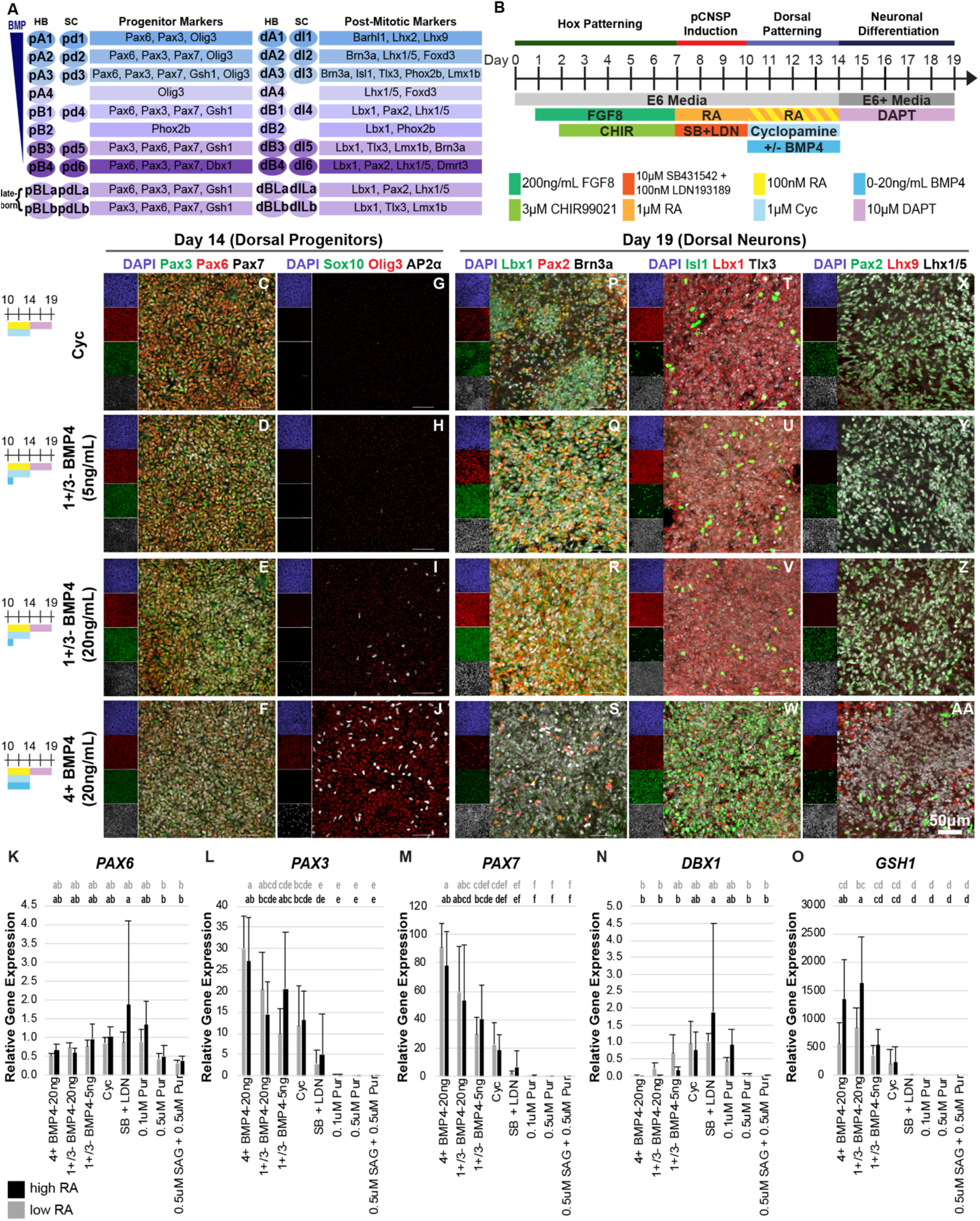
Concentration and Time Dependent BMP4 Patterning of Dorsal Spinal Neurons. (**A**) Dorsal pCNS populations, with characteristic progenitor and post-mitotic transcription factor markers for the hindbrain (HB) and spinal cord (SC). (**B**) Timeline of dorsal differentiation from H120-NMPs. (**C-J**) Immunostaining and (**K-O**) qRT-PCR in day 14 progenitor. Error bars represent standard deviation (n=6 biological replicates per condition). Data shown as relative gene expression compared to 100nM RA SB+LDN condition. Statistics were calculated by one-way ANOVA with Tukey-Kramer post-hoc (samples with the same letter assigned show no significant difference of p < 0.05). (**P-AA**) Immunostaining in day 19 post-mitotic neurons. All scale bars = 50μm. *See also Fig. S1 and S2*.

**Fig. 3.**
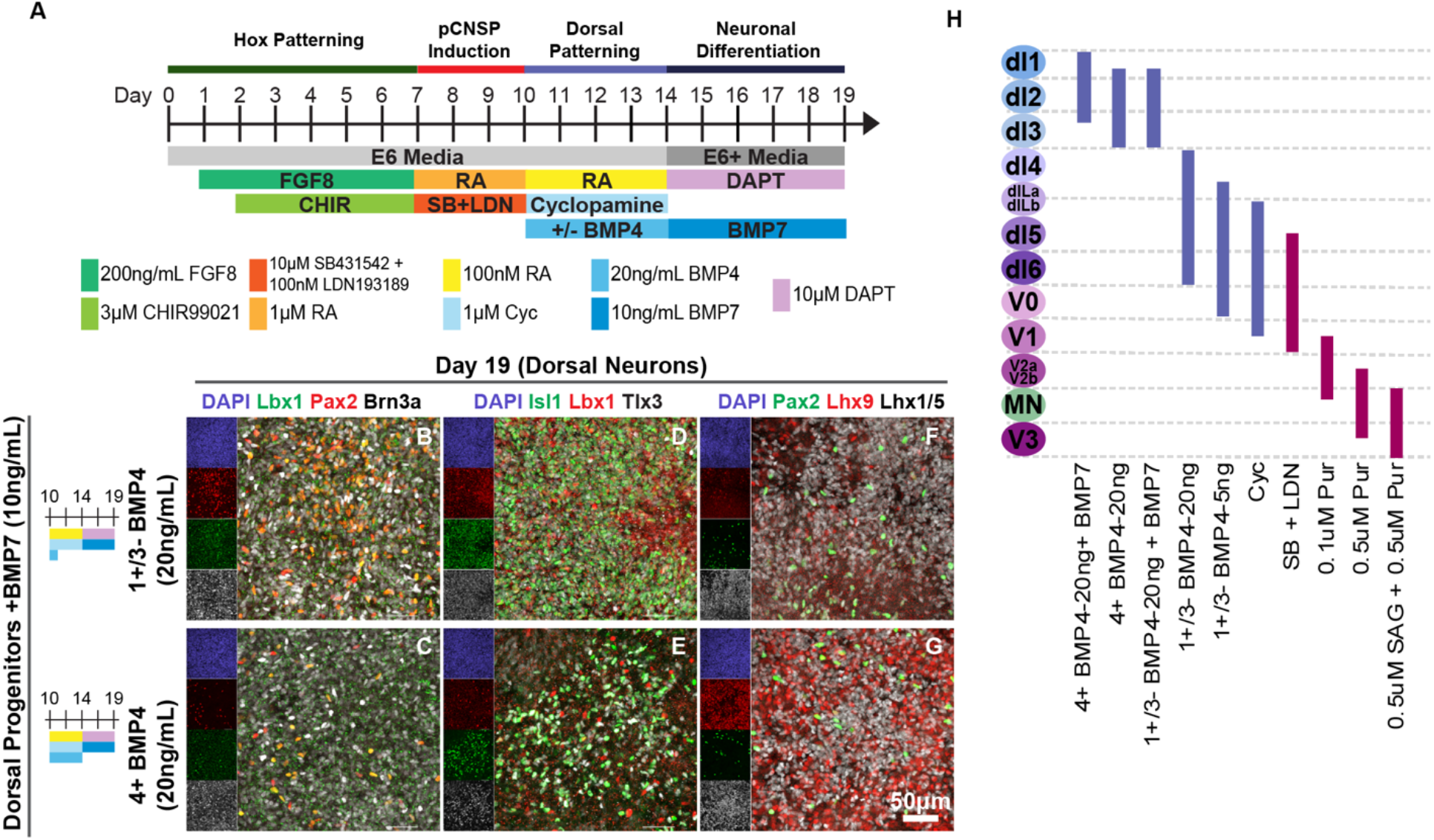
Addition of BMP7 During Neuronal Differentiation Further Dorsalizes Post-Mitotic Population. (**A**) Timeline of dorsal differentiation from H120-NMPs, with BMP7 added during neuronal differentiation phase from days 14-19. (**B-G**) Immunostaining in day 19 postmitotic cultures shows BMP7 ^+^ DAPT treatment rapidly converts progenitors to dorsally shifted post-mitotic phenotypes compared to Fig. 2. (**H**) Schematic of differentiation conditions corresponding post-mitotic cardinal cell types.

### Single Cell Transcriptomes Reveal Differential Population Distributions after Combined R/C and D/V Patterning

We generated an expansive scRNAseq dataset comprising dorsal and ventral populations differentiated from six NMP timepoints (H24, H48, H72, H120, H168, and H216) (**Fig. 4A,B, S3A-C**, **S4B**). For dorsal differentiation, pCNSPs were exposed to 100nM RA, Cyc, and pulsed with 20ng/mL BMP4 during the 4 day progenitor patterning period (**Fig. S3A,D**). For ventral differentiation, pCNSPs received 100nM RA, SB+LDN, 0.5uM and 0.5uM Pur (**Fig. S3B,E)**. Additionally in the D/V patterning stage, pCNSPs at H216 were patterned with either 1uM (H216R) or 100nM RA (H216) to determine whether RA further impacts caudalization (**Fig. S3A,B**). After DAPT treatment, the resulting samples were near homogenously neuronal (85-98% *SNAP25*^+^), with trace *SOX2*^+^ floor plate (*SHH*^+^) and roof plate (*LMX1A*^+^) cells and minimal expression of markers from other cell lineages, thereby demonstrating the efficiency of our modular differentiation methodology (**Fig. 4C**).

**Fig. 4.**
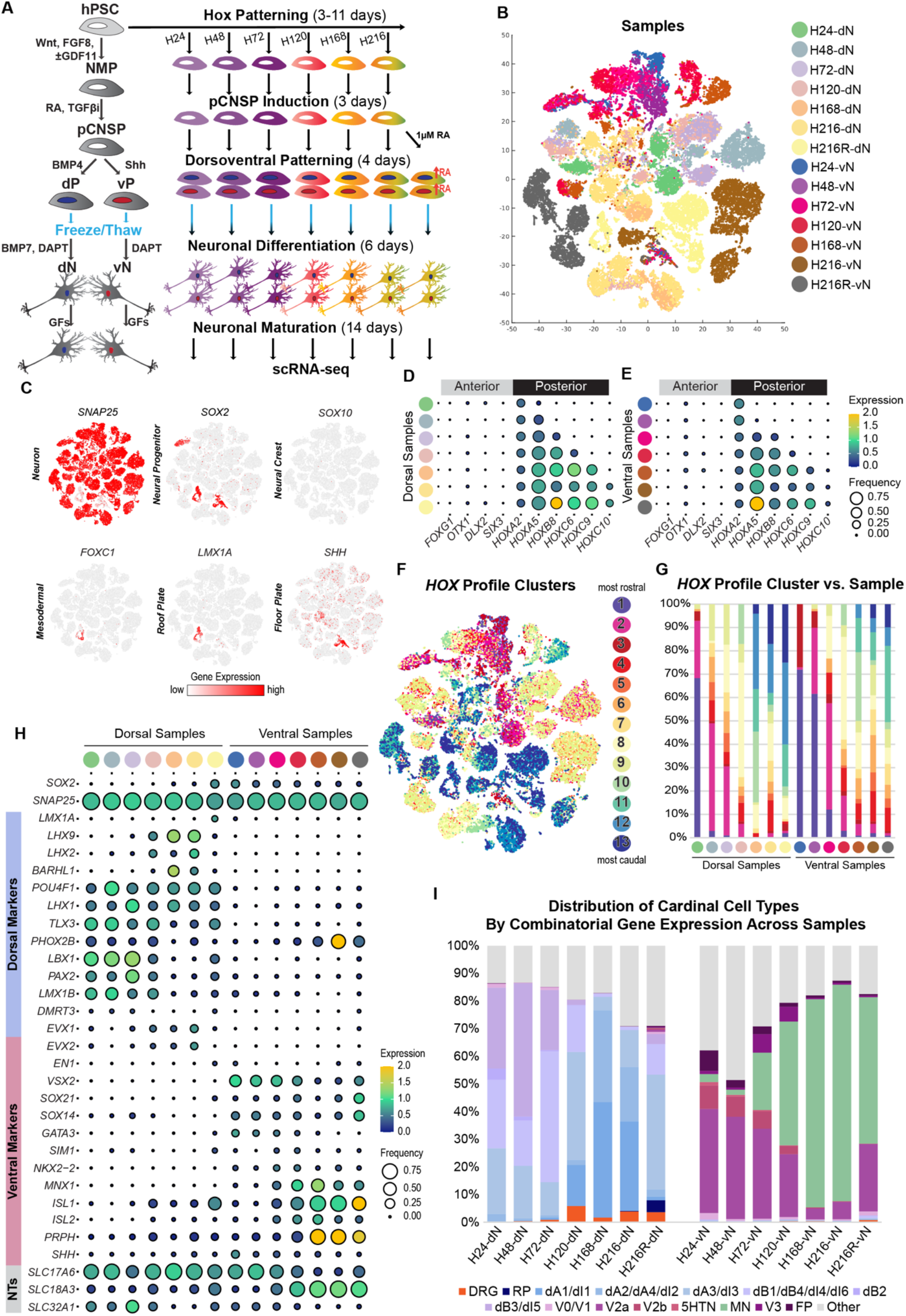
scRNAseq Characterization of Dorsal and Ventral Samples Differentiated from Discrete Regions Along R/C Axis. (**A**) Timeline of differentiation from region-specific NMPs, to discrete pCNSPs, dorsal (dp) and ventral (vp) progenitors, and post-mitotic neurons. (**B**) t-SNE plot with seven dorsal samples and seven ventral samples (n=46,959 cells). (**C**) t-SNE heatmaps showing highly neuronal (*SNAP25*^+^) cells, with few neural progenitor (*SOX2*^+^/*SNAP25*^-^), mesoderm (*FOXC1*^+^), or neural crest (*SOX10*^+^) cells. *SOX2*^+^ progenitors are primarily floor plate (fp; *SHH*^+^) and roof plate (rp; *LMX1A*^+^). (**D-E**) Dot plot displaying genes associated with anterior or pCNS identity across dorsal (**D**) and ventral (**E**) samples. The size of each circle reflects the fraction of cells where the gene is detected, and the color reflects the average expression level within each cluster (blue, low expression; yellow, high expression). (**F**) t-SNE plot showing *HOX* profile clusters. (**G**) Distribution of *HOX* profile clusters across samples. (**H**) Dot plot displaying genes associated with dorsal or ventral neuronal phenotypes. (**I**) Distribution of cardinal pCNS neurons, peripheral dorsal root ganglion (DRG) neurons, floor plate (FP), and roof plate (RP) cells as defined by non-overlapping combinatorial transcription factor expression across dorsal and ventral samples. *See also Fig. S1, S3 and S4*.

Analysis of *HOX* expression across all samples indicated discretization along the R/C axis, such that dorsal and ventral cultures derived from the same NMPs showed globally similar *HOX* expression (**Fig. 4D,E**). Compared to the previously presented scRNAseq dataset (**Fig. S1**), samples were rostrally shifted (**Fig. S4A,B**). This is likely a consequence of using SB+LDN during the pCNSP induction stage, which recapitulates the role of Noggin to abruptly terminate HOX progression in NMPs (Lamb and Harland, 1995; McMahon et al., 1998). Notably, increased RA concentration did not result in activation of more caudal *HOX* paralogs in H216R compared to H216 samples (**Fig. S4B**), confirming that R/C patterning occurs during NMP and pCNSP differentiation and is independent of RA concentration during D/V patterning. However, increased RA in H216R conditions did cause elevated expression of *HOXB8* and *HOXA5* in dorsal and ventral samples respectively (**Fig. 4D,E**, **S4B**), suggesting that RA may continue to play a role in neuronal subtype specification (Dasen and Jessell, 2009; Nolte and Krumlauf, 2013; Philippidou and Dasen, 2013).

We visualized simultaneous expression of all *HOX* genes by clustering, (**Methods, Fig. 4F,G**, **S4C**), which revealed inter- and intra-sample *HOX* profile heterogeneity, with caudal samples surprisingly inclusive of rostral *HOX* profile clusters (clusters 1-7) (**Fig. 4G**). These “mismatched” *HOX* profile clusters were associated with different phenotypes—including MNs, dI1, dI2, and dI3 INs (**Fig. 4F,G**, **5A,D**)—indicating neuronal subtype-specific *HOX* gene stratification in accordance with findings *in vivo* (Dasen and Jessell, 2009; Nolte and Krumlauf, 2013; Philippidou and Dasen, 2013). Thus, while *HOX* genes can be used globally to assess a sample’s R/C positional identity, nuances in *HOX* gene expression profiles of hPSC-derived neuronal subtypes also emerge with cell maturity and D/V specification. Our differentiation methodology may thus be used to explore how HOX dynamics influence pCNS neuronal specification and circuit organization (Philippidou and Dasen, 2013).

**Fig. 5.**
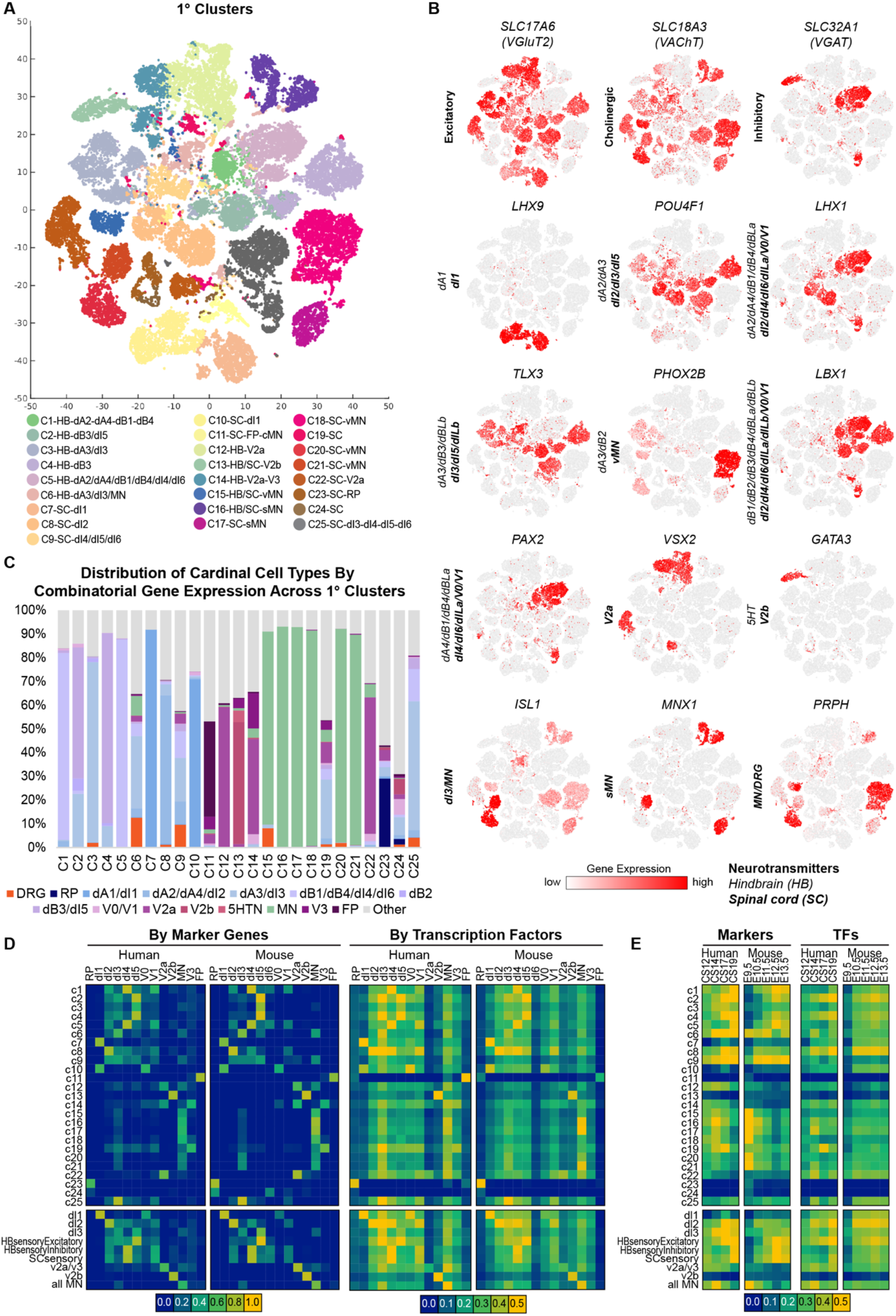
Unbiased NMF-based Clustering Validated by Global Similarities Between hPSC-derived and *In Vivo* scRNAseq Datasets. (**A**) t-SNE plot with 25 primary clusters broadly divides dataset by pCNS cardinal neuron identity. Legend labels indicate whether the cluster was presumed to be from the hindbrain (HB) or spinal cord (SC) (**B**) t-SNE heatmaps showing that characteristic cardinal transcription factors (hindbrain or spinal cord) and neurotransmitters (indicated by text legend) map to primary clusters. (**C**) Distribution of cardinal pCNS neurons, peripheral dorsal root ganglion (DRG) neurons, floor plate (FP), and roof plate (RP) cells as defined by non-overlapping combinatorial transcription factor expression across primary clusters. (**D-E**) Heatmap of Pearson’s correlation coefficient (PCC) values matrix comparing primary clusters or subpopulation groupings against *in vivo* human (Rayon et al., 2021) and mouse (Delile et al., 2019) embryonic (**D**) cardinal cell types and (**E**) developmental stages. Marker genes (n=77 for human; n=55 for mouse) were defined by the knowledge matrix provided in Rayon/Delile et al. and transcription factors (n=1,463 for human; n=1,775 for mouse) were defined by annotations from PANTHER and GO. *See also Fig. S5 and S6*.

We next assessed differentiation efficiency to various cardinal cell types. Dorsal and ventral samples showed gene expression patterns associated with appropriate transcriptional markers (**Fig. 4H**). The dataset was sparse in intermediate cardinal neurons corresponding to V0 (*EVX1*), V1 (*EN1*), and dI6 (*DMRT3*) INs, a consequence of using patterning conditions that yielded few DBX1^+^ progenitors (**Fig. S3D,E**). When we specifically examined cardinal cell type distributions defined by non-overlapping combinatorial transcription factor expression (**Table S1A**), we observed increasingly dorsal or ventral character as samples were caudalized. For example, dorsal H24-dN, H48-dN, and H72-dN samples corresponding to hindbrain-rostral cervical spinal cord were ventrally shifted toward *LBX1*^+^ dI4/dI5/dI6 INs compared to H120-dN, H168-dN, and H216-dN samples, which included primarily dI1/dI2/dI3 INs (**Fig. 4H,I**). Similarly, ventral H24-vN, H48-vN, and H72-vN samples were dorsally shifted to *CHX10*^+^ V2a and *GATA2/3*^+^ V2b INs compared to H120-vN, H168-vN, and H216-vN samples, which had a greater proportion of MNs (**Fig. 4H,I**). Given our application of a consistent D/V patterning protocol, this data suggests inherent differences in region-specific NMP differentiation potential. Moreover, though increased RA did not contribute to caudalization during progenitor patterning (**Fig. 4D,E, S4B**), higher RA exposure in H216R samples caused a shift toward more intermediate cell types compared to H216 samples (**Fig. 4I**), reaffirming our prior observation that RA modulates morphogen potency and is involved in neuronal fate determination.

### Unbiased Clustering Isolates Cardinal Cell Types

Although transcription factors that define pCNS cardinal cell types are greatly conserved during evolution (Marklund et al., 2014; Rayon et al., 2021), they could exclude potential species-specific or region-specific differences unique to our dataset. For example, if only cells expressing cardinal markers are analyzed, 15-50% of cells across our samples would remain uncharacterized (**Fig. 4I**, **Table S1B**). Therefore, we applied a clustering method based on sparse non-negative matrix factorization (NMF) (Kim and Park, 2008) to define 25 “primary clusters” (**Methods**, **Fig. 5A**). We assigned clusters to hindbrain or spinal cord based on sample identities’ global *HOX* expression (**Fig. 4D,E**) and assessed the composition of cardinal neurons in each cluster based on expression of known markers (**Fig. 5A-C**). To determine how our hPSC-derived clusters compared to *in vivo* neuronal populations, we performed a correlation analysis against recently available embryonic human (Rayon et al., 2021) and mouse (Delile et al., 2019) neural tube scRNAseq data across multiple gestational timepoints (**Fig. 5D, S5, Methods**). Both of these datasets relied on strict transcriptional definitions to define their cardinal neurons, a consequence of the sparsity of cells available for adequate clustering. Despite disparate approaches to cell type identification, we observed good similarity (Pearson Correlation Coefficient>0.5) between our clusters and the neuronal populations defined by Rayon et al. or Delile et al. using either known markers or their sets of annotated transcription factors (**Fig. 5D, S5**). A direct comparison between cells defined by known markers found similar concordance between *in vitro* and *in vivo* cell types as between the *in vivo* mouse and *in vivo* human cell types (Delile et al., 2019; Rayon et al., 2021)(**Fig. S5C**). Thus, we validated that our hPSC-derived populations were comparable to human and mouse neurons *in vivo*.

We then determined at what stage of *in vivo* development our hPSC-derived populations might belong by comparing our clusters to the Carnegie stage- (CS12-CS19) or mouse embryonic day-matched (E9.5-E13.5) samples from these studies (**Fig. 5E**). Dorsal clusters (C1-C10, C25) showed higher correlation to samples from CS17 and CS19 (gestational days 42-51), compared to ventral clusters (C11-C18, C20-23) which showed similarity to samples from CS12 and CS14 (gestational days 26-35). A comparable trend was observed in the mouse data. These correlation patterns were in accordance with the sequential emergence of ventral and dorsal neurons *in vivo*, wherein ventral populations are patterned earlier in development than dorsal populations (Marklund et al., 2014; Rayon et al., 2021). It is notable that our hPSC-derived neurons were derived in parallel and in fewer than 38 days, which suggests an accelerated differentiation of cells *in vitro* compared to endogenous populations. Altogether, these findings validate our data-driven approach to clustering pCNS neurons and present an opportunity to detect novel neuronal markers otherwise obscured by *a priori* transcriptional definitions or limited by the availability of embryonic tissue.

### Differential Gene Expression in Subpopulation Analysis of Primary Clusters

While many primary clusters were comprised of a single cardinal population, others were made up of closely related cell types. For example, C9 and C25 included multiple inhibitory and excitatory LBX1^+^ populations (dI4/dI5/dI6), and C14 contained both CHX10^+^ V2a INs and SIM1^+^ V3 INs, which are both glutamatergic ventral INs (**Fig. 5B-C**). We organized related primary clusters into 17 different groups (**Fig. 6A, Table S2A**), then developed and applied a consensus clustering based approach with the goal of defining robust subclusters representing subtypes of known cardinal populations (**Methods, Fig. S7A**). Consensus clustering was shown to improve cluster stability without sacrificing cluster quality (**Fig S7D, Methods**). Each subpopulation subdivided into 4-9 “subclusters”, to which we assigned a R/C positional identity based on sample identities’ global *HOX* expression (**Fig. 6B**, **Fig. 4D,E**). We examined the relatedness of these subclusters using hierarchical clustering, where subclusters generally organized by region (**Fig. 6B**). Analysis of differentially expressed genes (DEGs) across subclusters uncovered hundreds of genes upregulated in region-specific cardinal subtypes (**Table S3, Methods**). Here, we focus on the MN, dI1, and V2a/V3 groups, highlighting a few findings that emphasize how unbiased clustering enables the discovery of novel or region-specific markers different from or difficult to detect with available *in vivo* datasets. Similar analyses for other groups are available (**Table S3, <u>online resource</u>**).

**Fig. 6.**
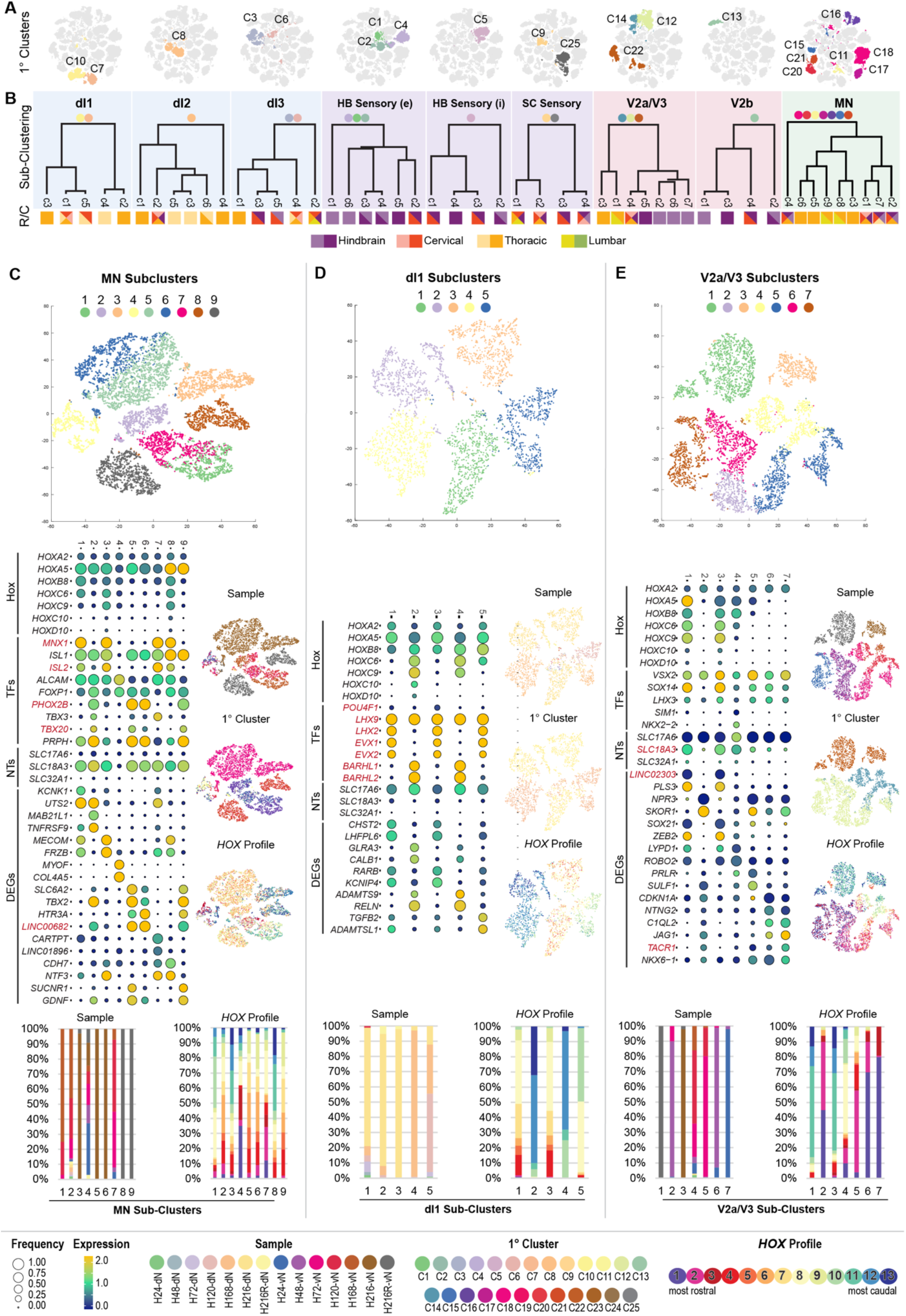
Subcluster Analysis Reveals Subpopulations with Neuronal Phenotype and Regional-Specificity. (**A**) t-SNE plots showing primary cluster compositions in 9 subpopulation groupings. (**B**) Hierarchal organization of subclusters with pictorial representations of estimated R/C location. Dual colors in key refer to rostral (pale) or caudal (dark) segments of hindbrain or spinal cord regions. (**C-E**) Subcluster analyses for (**C**) all MN populations, including sMN (C16, C17,C21), vMN (C18, C20, C21), and cranial MN (cMN; C11) clusters, (**D**) dI1 clusters (C7,C10), and (**E**) V2a/V3 mixed clusters (C12, C14, C22). t-SNEs show subclusters (n=5-9) defined by consensus clustering and distributions of the samples, primary clusters, and Hox profiles. Sample and *HOX* profile cluster distributions across subclusters are also visualized in stacked histograms. Dot plots display *HOX* gene expression, appropriate transcription factors (TFs) associated with the subpopulation grouping, neurotransmitters (NTs), and a selection or markers from the top 10 differentially expressed genes (DEGs) for each subcluster. The size of each circle reflects the fraction of cells where the gene is detected, and the color reflects the average expression level within each cluster (blue, low expression; yellow, high expression). *See also Fig. S7*.

MNs constitute the most widely studied neurons in the spinal cord, with significant evidence of Hox-dependent specification along the R/C axis and in the development of precise motor pools (Catela et al., 2016; Dasen and Jessell, 2009; Dasen et al., 2003, 2005, 2008; Liu et al., 2001; Mendelsohn et al., 2017; Philippidou and Dasen, 2013; Stifani, 2014). MNs in our dataset clustered into *MNX1*^+^/*ISL2*^+^ somatic MNs (sMNs; MN-c1,c3,c7,c8), which innervate skeletal muscle, and preganglionic *PHOX2B*^+^ visceral MNs (vMNs; MN-c2,c5,c6,c9), which are responsible for autonomic function (Guthrie, 2007; Stifani, 2014; Tiveron et al., 1996) (**Fig. 6C**). The latter population is a particularly rich target for novel findings. For example, Rayon et al. report scarce expression of *TBX20* as a notable difference between human and mouse vMNs, but hPSC-derived vMN subclusters in our dataset clearly express *TBX20*. *LINC00682* also emerged as a characteristic marker of vMNs in our dataset, but was upregulated in only p3 progenitors *in vivo* (Rayon et al., 2021). Though poorly understood, lincRNAs are abundant in the CNS and play multiple roles in development, neural plasticity, neurodegeneration, and sex-specific disease phenotypes (Issler et al., 2020; Quan et al., 2017; Wang et al., 2017). Given the origin of vMNs on the pMN/p3 border (Briscoe et al., 1999; Marklund et al., 2014), this could implicate *LINC00682* as an important gene regulator for vMN specification. Additionally, although sMNs and vMNs were proportionally divided within samples (**Fig. 4H, 6C**), *HOX* profiles appeared hindbrain-like in vMNs but sMNs maintained spinal *HOX* profiles correlative to their sample identity. *Phox2B* is known to be a direct target of several *Hox* genes (Philippidou and Dasen, 2013; Samad et al., 2004) and may contribute to this differential expression. Given that the persistence of Hox activity is thought to coincide with the development of downstream synaptic targets (Catela et al., 2016; Philippidou and Dasen, 2013), it is also possible that the proximity of ganglionic targets, both spatially and developmentally, causes early downregulation of *Hox* gene expression in vMNs compared to sMNs, a subject for future investigation.

The dI1 IN population is derived from the dorsal-most progenitor domain of the spinal cord (**Fig. 2A**) and migrates to the deep dorsal horn, where they have roles in proprioception (Bermingham et al., 2001; Helms and Johnson, 1998). DI1 subclusters divided into ipsilateral-projecting dI1i (*BARHL1/2*^+^; dI1-c2,c4) (Bulfone et al., 2000; Ding et al., 2012) and contralateral-projecting dI1c (*LHX2*^+^; dI1-c1,c3,c5)(Wilson et al., 2008) subtypes (**Fig. 6D**). The *LHX2*^+^ population also strongly expressed *EVX1/2*, which classically identify V0 INs (Marklund et al., 2014; Moran-Rivard et al., 2001; Pierani et al., 1999, 2001). In contrast to our dataset, *EVX1/2* are not expressed in dI1 INs in mouse or human scRNAseq data (Delile et al., 2019; Rayon et al., 2021). Furthermore, while hPSC-derived dI1 cells uniformly expressed *LHX9* (Helms and Johnson, 1998; Lee et al., 1998), they seldom co-express *POU4F1* (**Fig. 3F,G, 5B**), which is characteristic in mouse (Delile et al., 2019; Gowan et al., 2001; Lu et al., 2015) but may not be a consistent feature of the human population (Marklund et al., 2014; Rayon et al., 2021). *HOX* profiles appeared hindbrain-like in the *LHX2*^+^/*EVX1*^+^ population, but not the *BARHL1/2*^+^ population, which showed persistent caudal *HOX* profiles, despite comparable sample compositions (**Fig. 6D**). In particular, *Evx1* is regulated by *Hox2* paralogs (Davenne et al., 1999; Philippidou and Dasen, 2013), so its expression may suggest a potential role for *Hox* genes in gene regulatory pathways responsible for ipsilateral/contralateral projection patterns in dI1 neurons.

*CHX10*^+^ (*VSX2*^+^) V2a INs have multiple roles in locomotor coordination and breathing (Azim et al., 2014; Crone et al., 2008, 2012), and are one of the few spinal interneuron populations to have been characterized by spinal segment (Francius et al., 2013; Hayashi et al., 2018). Of the cardinal populations in our dataset, the V2a INs show the best continuous representation throughout all ventral samples (**Fig. 4I**). They also express *HOX* profiles commensurate with their sample identities (**Fig. 6E**). Region-specific markers associated with hindbrain (V2a/3-c2,c5,c6,c7) or spinal V2a INs (V2a/3-c1,c3) were apparent in subcluster DEGs, but these differed from markers identified with scRNAseq by Hayashi et al. This is likely because that dataset comprised FACS-sorted Chx10:tdTomato+ cells in p0 mouse cervical and lumbar tissues, which are developmentally advanced compared to our hPSC-derived cells (Hayashi et al., 2018). Spinal V2a INs in our dataset also expressed *SLC18A3* (*VACHT*) and *LINC02303*, which were not observed in more comparable mouse or human scRNAseq data (Delile et al., 2019; Rayon et al., 2021), though cholinergic character in V2a INs has previously been observed in zebrafish (Pedroni and Ampatzis, 2019). Finally, hindbrain V2a INs atypically expressed *NK1R* (*TACR1*) (**Fig. 6E**), which is normally expressed in pre-Botzinger complex (pre-BotC) respiratory neurons (Gray et al., 2001) and dorsal horn neurons (Sheahan et al., 2020; Todd, 2002, 2010). V2a INs in the rodent hindbrain are adjacent to the pre-BotC, but do not express NK1R (Crone et al., 2012), indicating a potential species-specific difference in rhythmic breathing organization.

Altogether, differences emergent in our hPSC-derived scRNAseq dataset reveal the power of this differentiation platform to generate novel, region-specific spinal subpopulations detectable by standard DEG analysis. Whether novel markers are bona fide, evidence of accelerated maturation of cells *in vitro* compared to *in vivo* (Dady et al., 2021), or artifacts of *in vitro* differentiation is subject to future investigation.

### Arboretum Analysis Reveals Complex Gene Expression Patterns Across Subclusters

While standard DEG analysis exposes strong differences between subclusters, it is restricted to pairwise comparisons. We reasoned that combinatorial patterns of gene expression could capture nuanced expression differences between subclusters and thus more comprehensively characterize novel cell types. To this end, we developed a computational pipeline that first applied Arboretum (Roy et al., 2013), a multi-task clustering algorithm, to cluster genes based on their expression states, calculated using pseudo-bulk mean gene expression. Then we identified “transitioning” gene sets, which demonstrate coordinated changes in expression states across subclusters (**Methods, Fig. 7A, <u>online resource</u>**). We interpreted these gene sets based on subclusters’ regional and phenotypic identities. Here we focus on the V2a/V3 and the high/low RA ventral groupings to demonstrate how this analysis can be used to detect patterns of interest in subpopulations along the R/C axis and to identify gene modules representing combinatorial expression changes across multiple cardinal populations in response to differentiation perturbations.

**Fig. 7.**
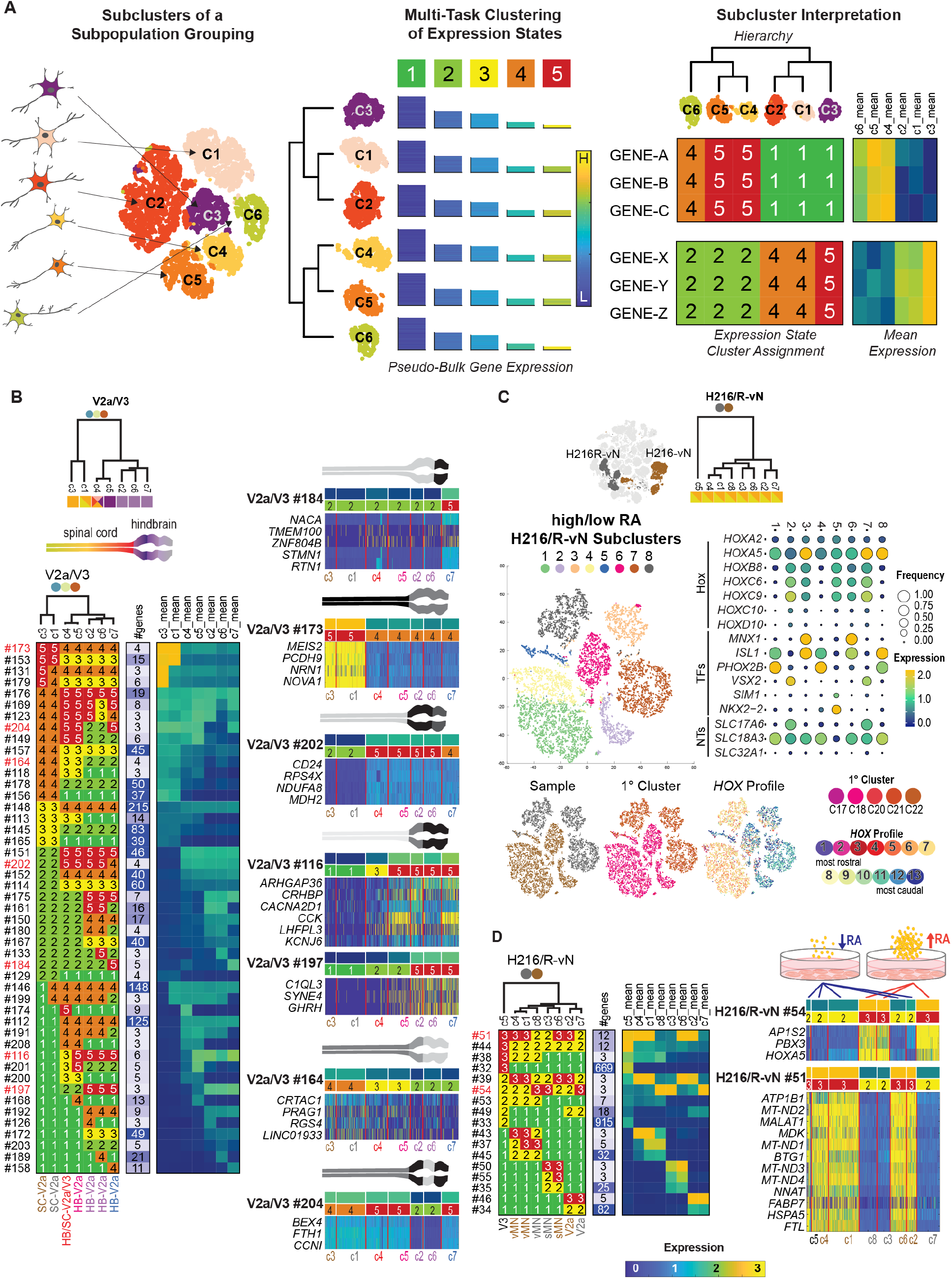
Multi-task Clustering Enables Discovery of Novel and Nuanced Gene Expression Patterns Across Subclusters. (**A**) Schematic procedure of Arboretum-based classification of DEG sets. Arboretum, a multi-task clustering method, clusters pseudo-bulk gene expression of each single cell subcluster with consideration of relationship structure between groups. Subsequent classification of genes is made by grouping genes that change their expression pattern across cell subclusters. (**B**) Expression state assignment patterns and mean expression levels for gene-sets identified for V2a/V3 subpopulation grouping. Subclusters colored by predominant primary cluster identity (top); R/C positioning (spinal cord (SC) or hindbrain (HB)) and cardinal cell type colored by sample identity (bottom). A selection of gene-sets highlight potential region-specific patterns of gene expression across the pCNS, represented in schematic images (right). (**C**) Subpopulation analyses for grouping of ventral samples exposed to high (H216R-vN) or low (H216-vN) RA. TSNE plots show subclusters (n=5-9) defined by consensus clustering and distributions of the samples (full dataset and high/low RA subpopulation) and *HOX* profiles. Hierarchal organization of subpopulation subclusters with pictorial representation of comparable thoracolumbar identity. (**D**) Expression state assignment patterns and mean expression levels for gene-sets identified for high/low ventral subpopulation grouping. Subclusters (top) and cardinal cell type composition (bottom) colored by sample identity. A selection of gene-sets highlights shared genes across multiple cardinal populations upregulated in response to high or low RA (right)

Because the V2a/V3 grouping divided into subclusters corresponding to R/C regions from the hindbrain through thoracolumbar spinal cord, the transitioning genes from Arboretum indicated region-specific expression patterns (**Fig. 7B**). Akin to DEG analysis, we found patterns of expression specific to a single subcluster, like V2a/3-#184, which shows elevated expression in V2a INs (c7) of the rostral hindbrain. We also found shared patterns of expression between multiple subclusters, like V2a/3-#173 or #202, which show genes upregulated in spinal or hindbrain subtypes respectively. These gene sets included factors of potential interest involved in binding HOX proteins (*MEIS2*), mitochondrial activity (*RPS4X, NDUFA8, MDH2*), cell adhesion (*PCDH9*), surface biomarkers (*CD24*), neurite outgrowth (*NRN1*), and neuron-specific alternative splicing (*NOVA1*). We also found patterns representing gradual changes in gene expression that may identify region-specific changes that emerge as a gradient along the R/C axis or between subclusters, as has previously been observed for V2a INs (Hayashi et al., 2018). For example, V2a/V3-#116 and #197 show gradual decrease in gene expression from hindbrain to spinal V2a INs, while V2a/V3- #164 shows gradual increase in expression from hindbrain to spinal V2a INs. Finally, we identified numerous gene sets that correspond to nuanced gene expression patterns, like V2a/V3-#204, which exhibit high levels of gene expression in the spinal cord and rostral hindbrain, but not the caudal hindbrain. The Arboretum pipeline and associated analysis is thus a valuable resource for curation of novel gene expression patterns that can be examined with targeted *in vivo* studies, compared to standard DEG analysis which fails to contextualize or detect nuanced gene expression differences between subclusters.

Next, we used Arboretum to determine whether changing RA during D/V patterning had an impact on terminal gene expression. We observed that changing RA concentration during dorsal differentiation significantly changed the distribution of post-mitotic cardinal populations (**Fig. 4I)**, but while ventral populations were slightly shifted, both H216-vN and H216R-vN samples contained V2a INs, sMNs, and vMNs **(Fig. 7C**). This allowed for direct comparison between cardinal populations. Arboretum identified gene sets comprising commonly upregulated (H216/R-vN-#54) or downregulated (H216/R-vN-#51) genes in response to the increase in RA concentration **(Fig. 7D**). Gene set #54 includes *HOXA5*, which validates the role of RA in activation of specific Hox genes and mimics occurrences *in vivo*. Constitutive activation of RA signaling during development was found to disrupt digit-innervating MN development (Mendelsohn et al., 2017), and precise retinoid levels are required for digit and tendon development (Rodriguez-Guzman et al., 2007). Notably, the annotation of genes in H216/R-vN- #51 (**<u>online resource</u>**) indicated that ventral patterning with 1μM RA compared to 100nM RA persistently suppressed signaling pathways involved in mitochondrial electron transport, mitochondrial respiration, and oxidation-reduction in post-mitotic neurons matured 20 days beyond progenitor patterning. This finding could have significant implications for *in vitro* modeling of neurological disorders associated with mitochondrial pathologies and cell survival after transplantation. Moreover, it highlights the need for more thorough characterization of differentiation protocols used for prospective cell therapies, which may optimize for a particular cell phenotype without considering how subtle changes in morphogen concentrations can impact long-term transplant efficacy.

## Discussion

By implementing a modular differentiation paradigm that explicitly decouples R/C from D/V patterning, we demonstrate the ability to direct hPSCs to any neuronal phenotype in the pCNS. Importantly, we show that all D/V phenotypes--particularly dorsal INs—can be effectively generated in monolayer culture conditions. This is in contrast to the previously requisite organoid and spheroid cultures, which exhibit batch-to-batch variability and may rely on the formation of signaling centers for D/V patterning (Andersen et al., 2020; Duval et al., 2019; Gupta et al., 2018; Ogura et al., 2018; Zheng et al., 2019). Moreover, our patterning schema enables deeper investigation of the role of Hox genes, retinoids, and other signaling molecules in the development of anatomically and therapeutically relevant cell types.

Our scRNAseq data also highlights the power of hPSCs to provide broad access to embryonic pCNS tissues. While scRNAseq datasets from primary embryonic rodent (Delile et al., 2019; Russ et al., 2021) and human spinal cords (Rayon et al., 2021) are invaluable resources, they have limitations. Because of the physical difficulties associated with early embryonic tissue acquisition and dissection, these datasets fail to discretize neuronal phenotypes across different pCNS R/C regions. They also sparsely sample individual cell subtypes, a consequence of poor neuronal yield and subtype rarity. By comparison, our modular protocol enables unlimited sampling of any phenotype from any differentiation timepoint across any R/C region. As a result, our scRNAseq dataset spans multiple discrete regions from the hindbrain through thoracolumbar spinal cord, improving the ability to detect nuanced transcriptional programs that potentially regulate lineage specification.

The multi-regional nature of our dataset posed challenges to systematically define cell clusters using existing methods. While known cell markers are used commonly to define single-cell populations (Delile et al., 2019; Rayon et al., 2021), a large proportion of cells remain unlabeled. In contrast, a clustering-based approach offers a more comprehensive strategy but remains challenging especially when there are a large number of unknown populations (Kiselev et al., 2019). Our two-step approach based on sparse non-negative matrix factorization and consensus clustering allowed a biologically meaningful grouping of cells that recapitulated known as well as novel cell types. Furthermore, our Arboretum-based approach allowed us to uncover previously unknown patterns of expression that can inform functional experiments for in-depth characterization of these cell populations. We anticipate that these platforms will enable rigorous interrogation of gene regulatory pathways responsible for neuron diversification and synaptic targeting in the pCNS. Future studies encompassing other pCNS populations including those from patient-iPSCs will enable investigation of spatiotemporal gene expression dynamics during development and disease.

## Acknowledgements

The authors thank the University of Wisconsin-Madison Biotechnology Center Gene Expression Center & DNA Sequencing Facility for providing single nuclei library preparation and next generation sequencing services. Single cell RNAseq data was analyzed by the UW Bioinformatics Resource Center. We also thank C. Birchmeier and T. Müller for the generous gifts of Lbx1, Tlx3, and Lmx1b antibodies. We thank N. Fedorchak for technical and editing assistance.

This work was made possible by US EPA grant 83573701 (RSA and SR), NIH NIGMS grant 1R01GM117339 (SR), NIH NCATS grant UG3TR003150 (RSA), NSF CAREER Award #1651645 (RSA), University of Wisconsin-Madison startup funds (RSA and SR), an Innovation in Regulatory Science Award from BWF (RSA), and a UW Data science foundation grant (SR). N.I. was supported by postdoctoral fellowships from the University of Wisconsin Stem Cell and Regenerative Medicine Center (SCRMC) and NIH-NINDS (F32 NS106740). JS was supported by a James McDonell Foundation Grant (#3194-133-349500-4-AAB5159).

## Author Contributions

Conceptualization, N.I., R.S.A., and S.R.; Methodology, N.I., J.S., R.S.A., and S.R.; Software, J.S. and S.R.; Formal Analysis, N.I. and J.S.; Investigation, N.I., J.S., S.C., Y.T., T.D., N.N., S.G.M.; Resources, N.I, R.S.A., and S.R.; Data Curation, N.I, J.S, S.R,, and R.S.A.; Writing – Original Draft, N.I., J.S., R.S.A., and S.R.; Writing – Review & Editing, N.I., J.S., R.S.A., and S.R.; Supervision, R.S.A., and S.R.; Project Administration, R.S.A. and S.R.

## Supplementary Figures

**Fig. S1.**
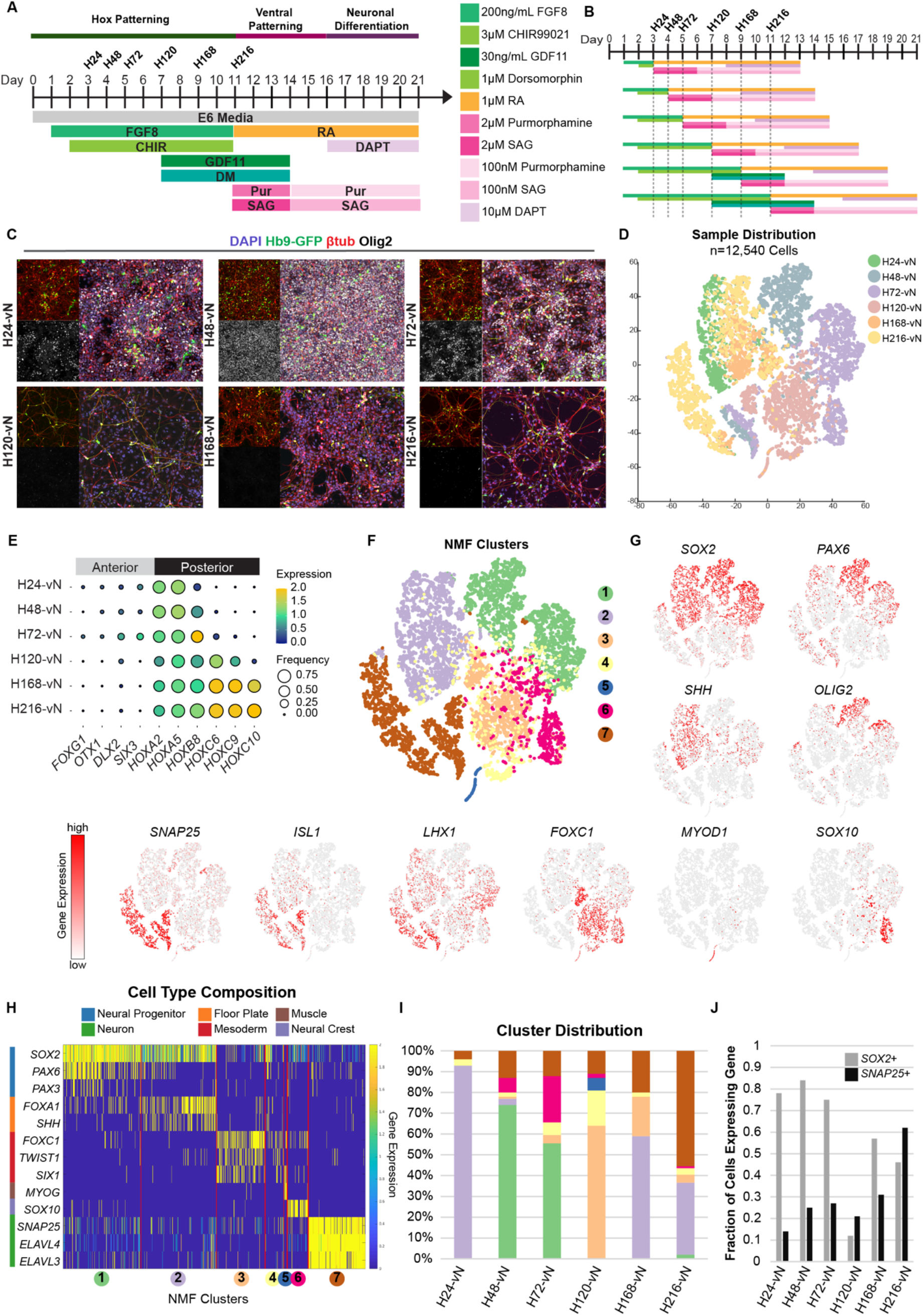
Direct Differentiation of pCNS Neurons from NMPs (Related to Fig. 1 and 4) (**A,B**) Timeline of HUES3-Hb9-GFP differentiation adapted from (Lippmann et al., 2014). Six neuronal samples were generated for analysis and scRNAseq, corresponding to 24hrs (H24), 48hrs (H48), 72hrs (H72), 120hrs (H120), 168hrs (H168), or 216hrs (H216) of Hox patterning. NMPs were patterned with Shh agonists SAG and Pur for 5 days, then treated with DAPT for 5 days prior to cryopreservation. Samples were thawed overnight prior to analyses. (**C**) Immunostaining of thawed cultures for GFP (green, MNs), β-tubullin III (red, neuronal), Olig2 (white, pMN), and nuclei (blue, DAPI). Scale bar = 50μm. (**D**) TSNE plot with six ventral samples (n=12,540 cells). (**E**) Dot plot displaying genes associated with anterior or pCNS identity across samples. The size of each circle reflects the fraction of cells where the gene is detected, and the color reflects the average expression level within each cluster (blue, low expression; yellow, high expression). (**F**) TSNE plot with seven determined clusters. (**G**) TSNE plots and (**H**) heatmap showing gene expression levels of characteristic markers for 6 major populations: neural progenitors (*SOX2, PAX6, OLIG2*), neurons (*SNAP25, ELAVL3, ELAVL4, ISL1, LHX1*), floor plate (*SHH, FOXA1*) mesoderm (*FOXC1, TWIST1, SIX1*), muscle (*MYOD1, MYOG*), and neural crest (*SOX10*). (**I**) Distribution of clusters across samples demonstrating cell type heterogeneity. (**J**) Fraction of *SOX2*^+^ neural progenitors and *SNAP25*^+^ neurons across samples.

**Fig. S2.**
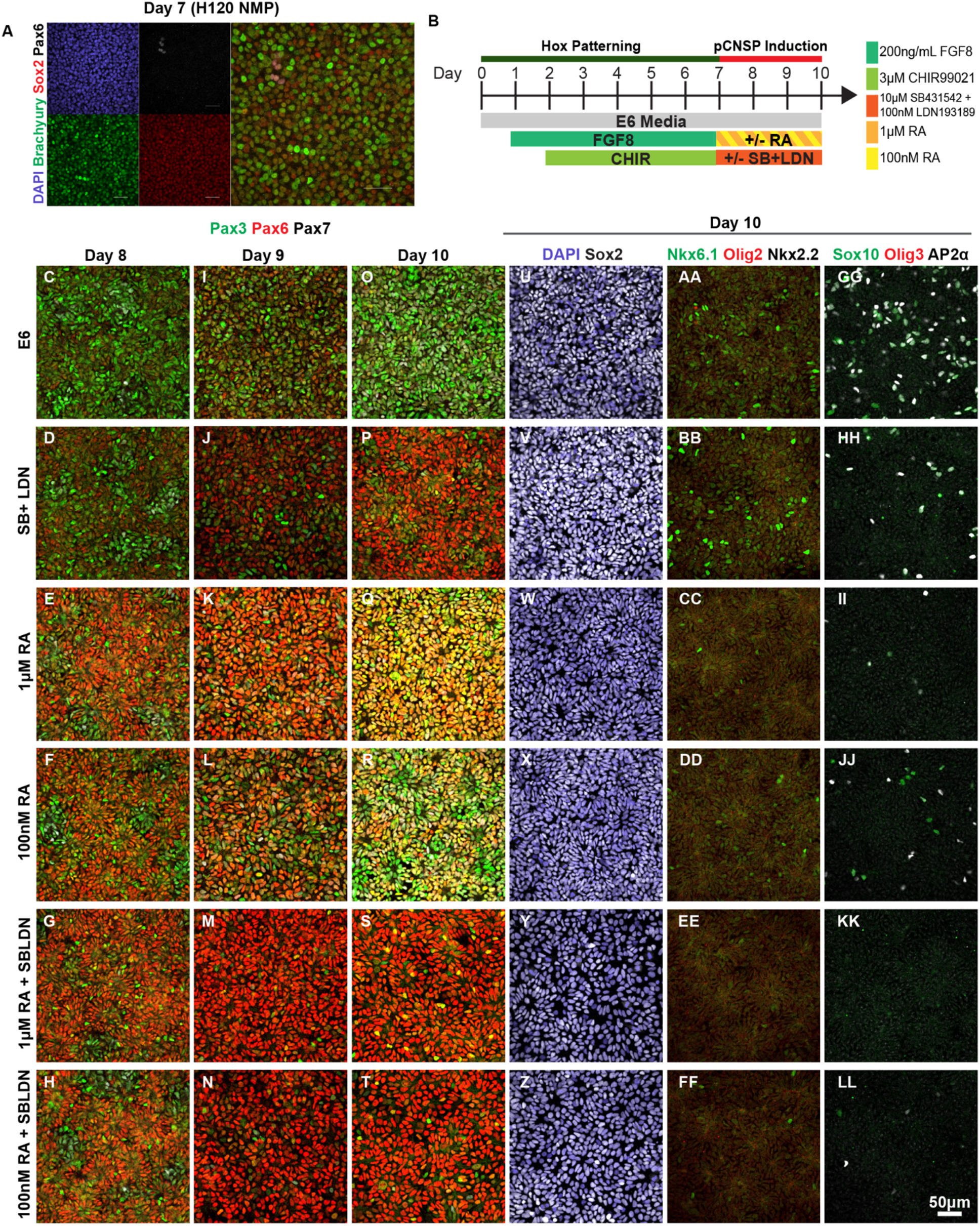
RA and Dual SMAD Inhibitors Cooperate to Efficiently Convert NMPs to Naive pCNSPs (Related to Fig. 1 and 2) (**A**) Immunostaining in H120-NMPs showing DAPI^+^ nuclei co-expressing BRACHYURY/SOX2 but not neuroectoderm marker PAX6. (**B**) Timeline of induction from H120-NMPs to pCNSPs. (**C-T**) Immunostaining between days 8-10 showing expression of PAX3, PAX6, and PAX7. (U-LL) Immunostaining on day 10 showing maintenance of SOX2^+^ neural progenitors (**U-Z**) and presence or absence of ventral (**AA-FF**), dorsal and neural crest markers (**GG-LL**). All scale bars = 50μm.

**Fig. S3.**
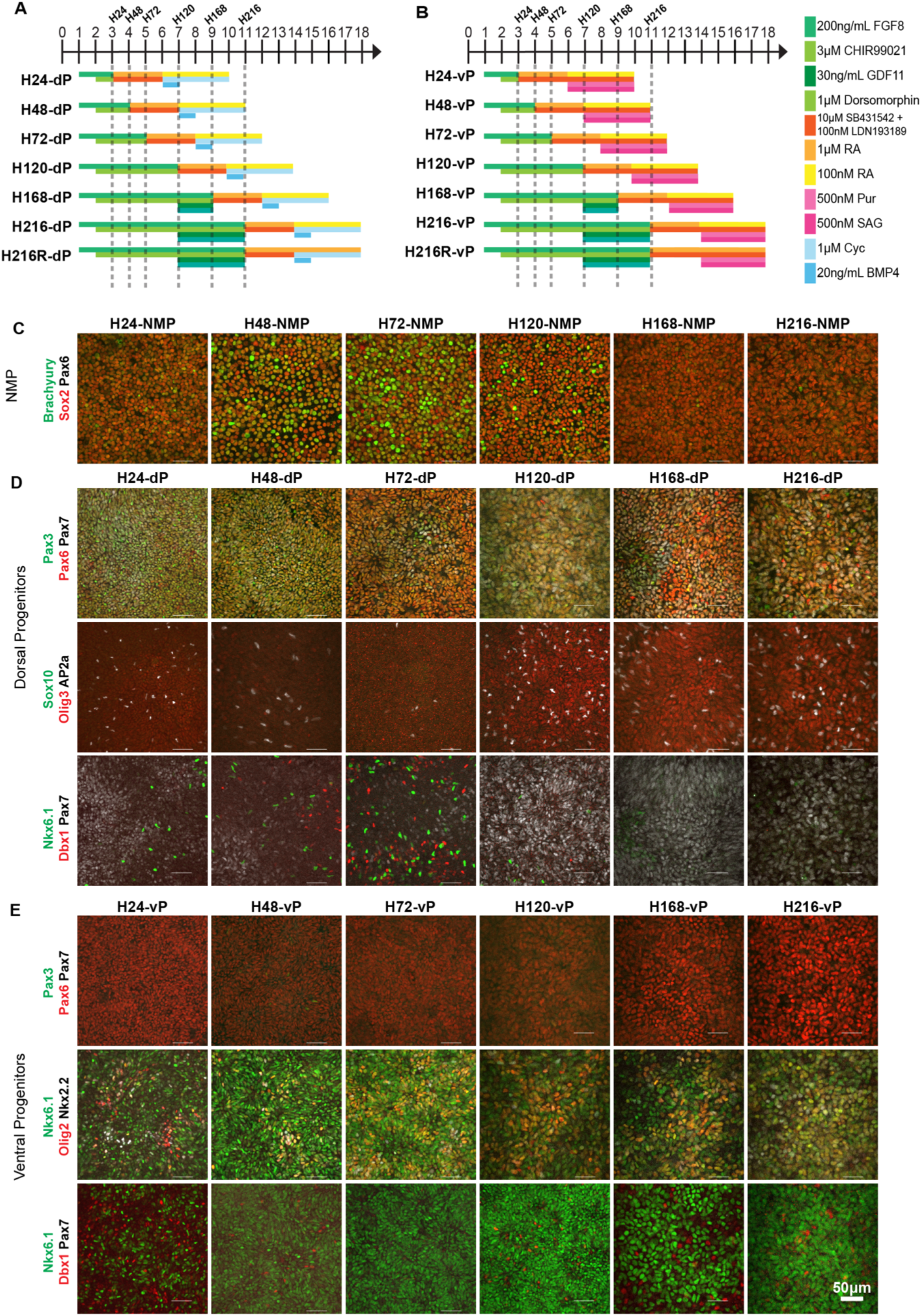
Generation of Dorsal and Ventral Progenitors from Multiple NMP Populations along R/C Axis (Related to Fig. 4) (**A,B**) Timeline of dorsal progenitor (dP) (**A**) and ventral progenitor (vP) (**B**) differentiation from NMP timepoints corresponding to 24hrs (H24), 48hrs (H48), 72hrs (H72), 120hrs (H120), 168hrs (H168), or 216hrs (H216) of Hox patterning. “H216R” conditions replace 100nM RA with 1uM RA during progenitor patterning with dorsal or ventral morphogens. Progenitors were cryopreserved after differentiation to synchronize cultures prior to maturation and scRNAseq. (**C**) Immunostaining in NMPs showing co-expression of BRACHYURY (green) and SOX2 (red), with the absence of PAX6 (white, spinal progenitor). (**D-E**) Immunostaining in dorsal (**D**) and (**E**) ventral progenitor cultures with characteristic markers.

**Fig. S4.**
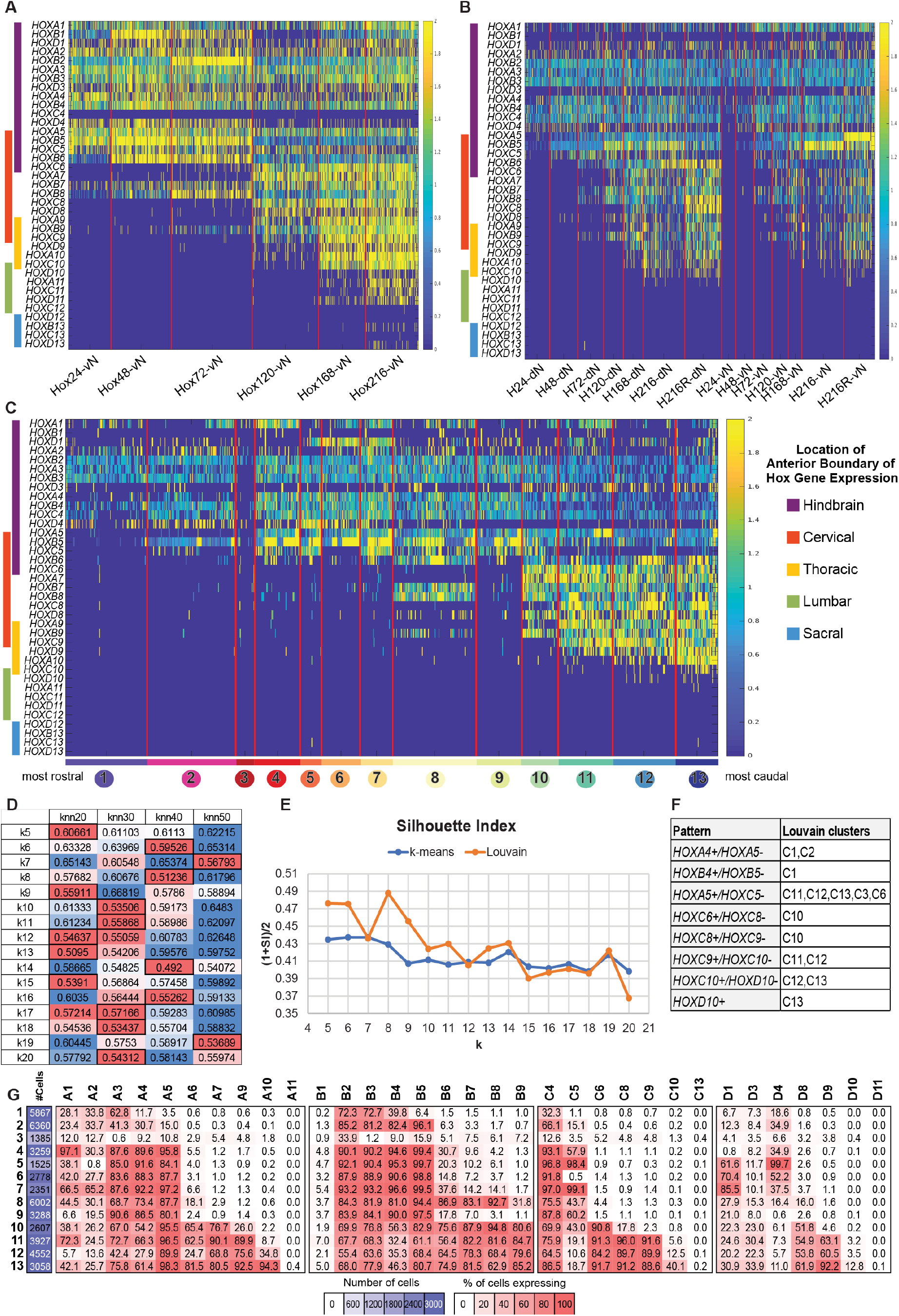
HOX Gene Expression Profiles for scRNAseq Datasets (Related to Fig. 4 and 6) Heatmaps showing expression levels of 33 *HOX* genes (**A**) across samples in the pre-optimized dataset in Fig. S1, (**B**) across samples in the optimized dataset in Fig. 4, and (**C**) across Hox profile clusters in the optimized dataset in Fig. 4G. (D) Heatmap of Pearson’s correlation values used for the selection of optimal number of kNN graph neighbors and number (columns) of cluster (rows). More reddish cell means containing more distinctive *HOX* profiles. Final selections are highlighted by outlines. (E) Silhouette indices (SI) for each k numbers of Louvain clustering and the k-means clusters. (F) Captured known *HOX* colinear expression patterns from the selected *HOX* profile Louvain cluster k=13. (G) Patterns were identified by observing the heatmap of clusters’ composition for *HOX* gene expressing cells. Captured patterns of other k numbers are available in Table S5.

**Fig. S5.**
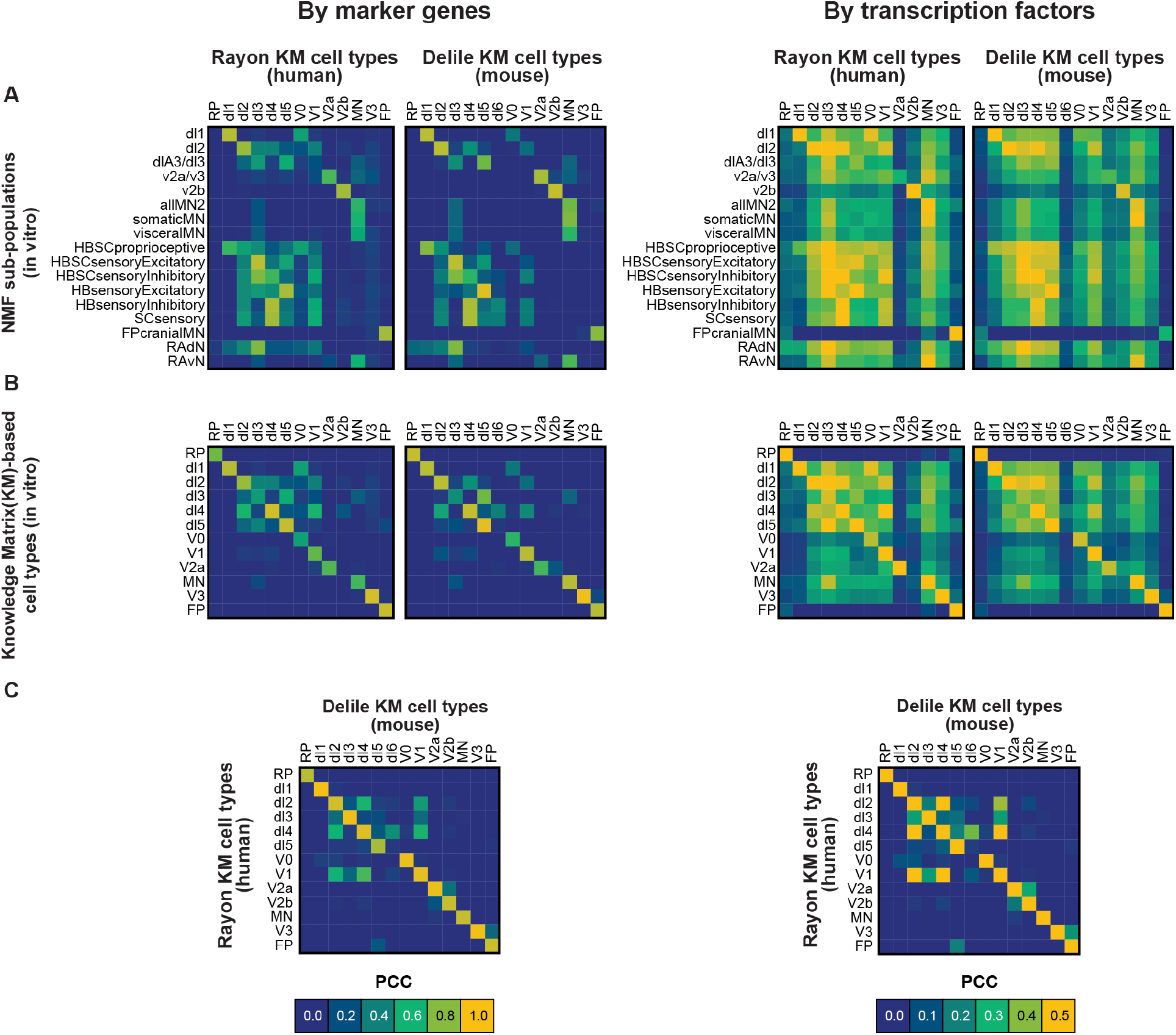
Pearson’s Correlation Coefficient Matrices for hPSC-Derived and *In Vivo* scRNA-seq Datasets (Related to Fig. 5) Heatmaps of Pearson’s correlation coefficient (PCC) values matrix. (**A**) 17 subtypes of primary and (**B**) 12 cell types defined by knowledge matrix are compared against in vivo human (Rayon et al.) and mouse (Delile et al.) embryonic cardinal cell types, using marker genes (n=77 for human; n=55 for mouse) defined by the knowledge matrix provided in Rayon/Delile et al. and transcription factors (n=1,463 for human; n=1,775 for mouse) defined by the annotations from PANTHER and GO. (**C**) Comparison of in vivo human (Rayon et al.) and mouse (Delile et al.) cardinal cell types.

**Fig. S6.**
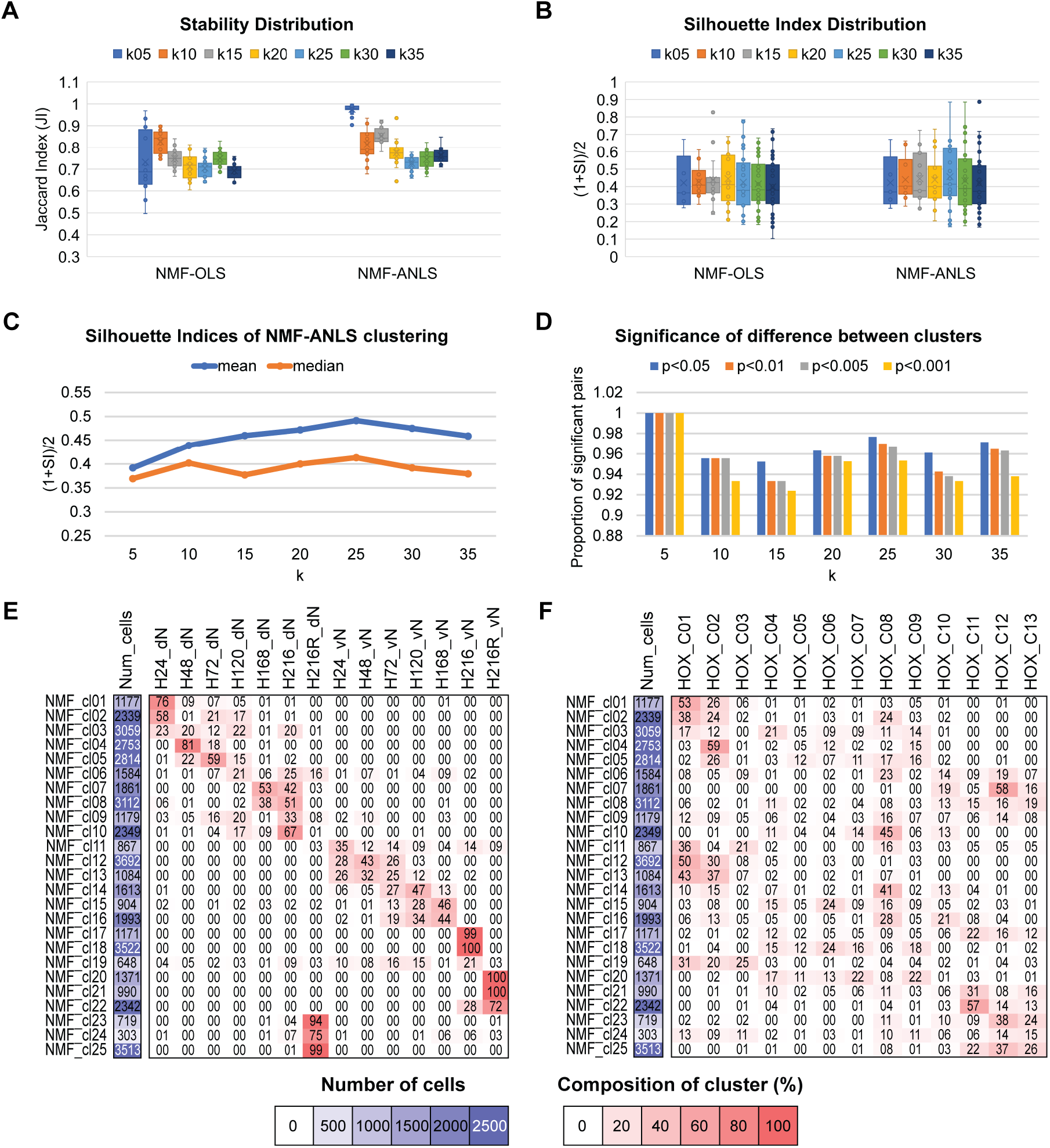
Specification of Optimal Parameters in the NMF-Based Identification of Primary Clusters (Related to Fig. 5) Specification of optimal parameters in the NMF-based identification of primary cardinal cell types. (**A**) Distribution of cluster-wise silhouette indices (SI) per k numbers from two different NMF algorithms: NMF-OLS and NMF-ANLS. (**B**) Distribution of Jaccard indices (JI) among 20 different random initializations per k numbers from two different NMF algorithms. (C) SI values of different k numbers of NMF-ANLS calculated from the profiles made of 23 well known neural cell type marker genes. (**D**) Counted proportions of combinatorial pairwise clusters that have shown significant differences in expression profiles. (**E**) Heat maps of clusters’ compositions of cells per 14 samples. (**F**) Heat maps of clusters’ compositions of cells per 13 *HOX* profiles clusters.

**Fig. S7.**
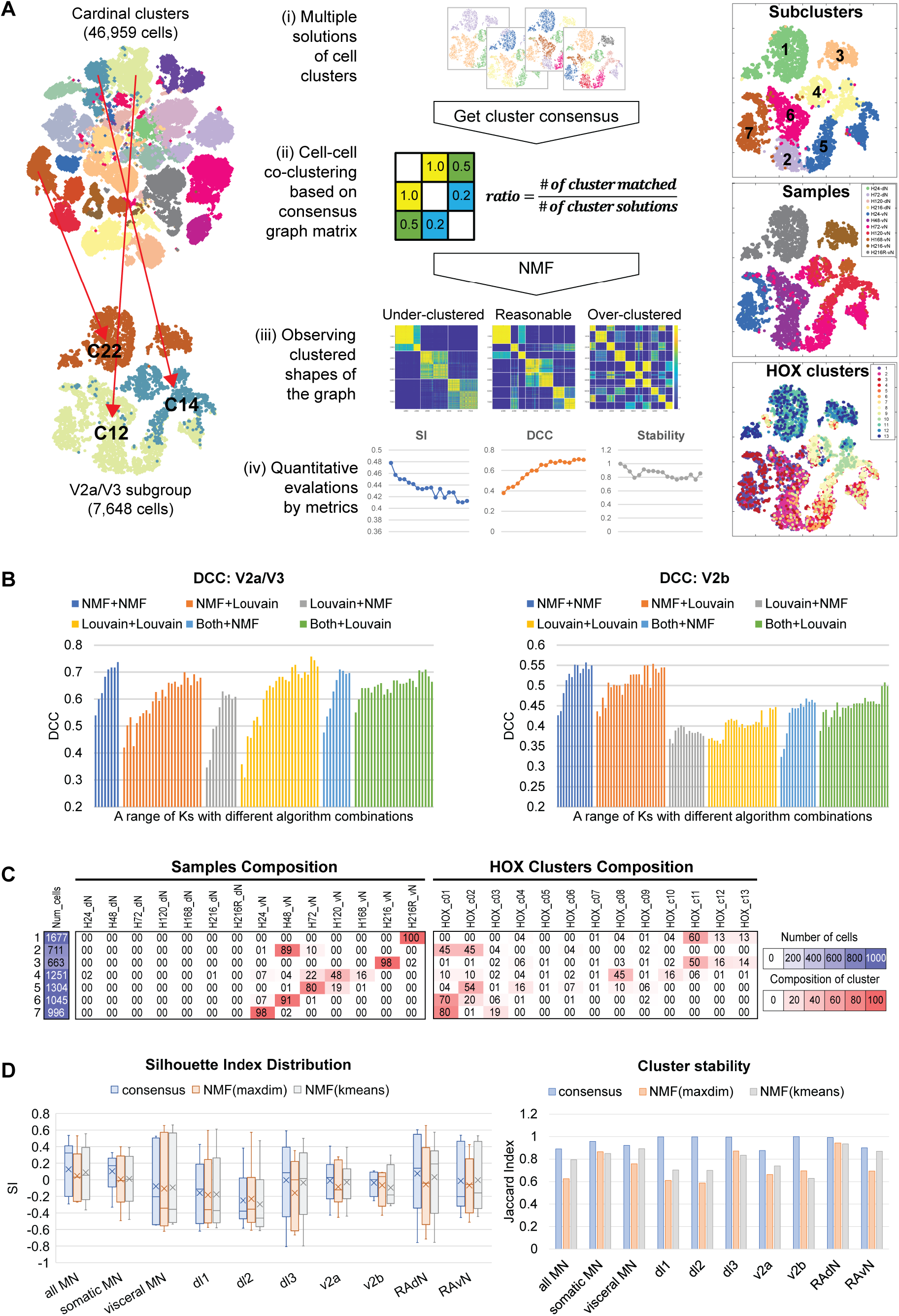
Identification of Subclusters Based on Consensus Clustering (Related to Fig. 6) (**A**) Schematic procedure of subpopulation clustering analysis based on the consensus clustering. 17 different neuronal groups were taken by partial regrouping of 25 primary clusters. Shown example is for the V2a/V3 group (left). Each group has been clustered based on several different k numbers and clustering methods to generate multiple solutions of cell clusters (middle, i). Then, multiple clustering results were summarized as a ratio value which enumerates the co-occurrence between cells within the same cluster (middle, ii), which results in a symmetric [cells x cells] consensus graph of all cells in the group. Finally, we clustered the consensus graph matrix to find the robust cell clusters or subtypes. For the specification of best k number, we used manual inspection of the block-diagonalness of the clustered consensus graph matrix to avoid over- or under-clustering (middle, iii), as well as evaluation metrics including silhouette index (SI), delta consensus counts (DCC) and stability (middle, iv). After we obtained the robust subclusters we compared the subcluster’s composition of time-dependent samples and *HOX* profiles clusters to annotate the regional identities (right). (**B**) Evaluation of different clustering algorithms on consensus clustering based on DCC. Our consensus clustering consists of two types of clusterings: clustering for the generation of ensemble and clustering of the consensus graph matrix. For the generation of the ensemble of clusters, we tested 3 different types of algorithms, which were NMF (“NMF”), kNN-based Louvain clustering (“Louvain”), and using all result from both algorithms (“Both”). For the consensus clustering, we tested 2 different types of algorithms, which were NMF of the graph matrix (“NMF”) and graph Louvain clustering (“Louvain”). 6 different combinations of algorithms of “ensemble generation + consensus clustering”, such as “NMF+NMF” or “Louvain+NMF”, are depicted in different colors of bar plots. We test several numbers of ks which correspond to each narrow bar. (**C**) Samples (left) and *HOX* profiles (right) compositions of 7 subpopulations identified from the consensus clustering of V2a/V3 group. (**D**) Comparison performances between the consensus clustering method and single NMF clustering by taking maximal factors (NMF maxdim) or single NMF clustering by performing additional k-means upon the factorized matrix (NMF kmeans). Performances are evaluated by silhouette index distribution (left) for the clustering quality and by clustering stability from 10 different random initialization (right).

## Supplementary Table Legends

**Table S1.** Knowledge matrix and cardinal cell type distribution (Related to Fig. 4 and 5) (A-B) Cardinal cell type distributions of samples (A) or 25 primary clusters (B) defined by non-overlapping combinatorial expression of 23 transcription factors. (C) Knowledge matrix of cell types and 23 defining transcription factors curated from literature.

**Table S2.** Organization of 17 neuronal groups and their subclusters (Related to Fig. 5 and 6) 17 neuronal groups organized from the 25 primary clusters. Constituent primary clusters or samples are listed in column B. 4-9 subclusters were identified by consensus clustering for each group. Approximated regional identities are listed in column D-M.

**Table S3.** Differentially expressed genes (DEGs) of 4-9 subclusters of 17 neuronal groups identified from conventional statistical methods (Related to Fig. 6) (A) Stat table of counts of DEGs per subtype. (B-R) 17 neuronal group-wise DEG lists with evaluation ratios and statistical p-values. Each list consists of 8 columns which designates subcluster ID (column A), gene ID (column B), rank in each subcluster (column C), ratio number of expressing cells within-cluster/outside-of-cluster (column D), log2 ratio of mean expression values within-cluster/outside-of-cluster (column E), hypergeometric p-value (column F), T-test p-value (column G), and MWW-test p-value (column H). Note that only genes that are expressing more than 50% of the subtype cells were counted as DEGs.

**Table S4.** Antibodies and Taqman Primers (Related to Fig. 1-3) (A) Primary antibody vendors and dilutions. (B) Taqman primer information for qRT-PCR.

**Table S5.** Observed specific expression patterns of HOX profile clusters used in the specification of k number for HOX profiles Louvain clustering. (Related to Fig. 4) 8 representative *HOX* expression patterns and the cluster IDs which have shown correspondence to the pattern for k=5-20 clusters generated by the *HOX* profiles Louvain clustering.

**Table S6.** Parameters used in the specification of subclusters and Arboretum analysis (Related to Fig. 6 and 7) (**A**) K number specification for the number of subclusters per 17 neuronal groups. Calculated evaluation metrics are designated. Bold fonts are top 3-5 candidates and red fonts are the final decision which is supported by manual inspection of block-diagonalness of the consensus graph matrix. (**B**) Distance metrics used in the generation of hierarchical trees of 17 neuronal groups (column C). We used unweighted average distance (UPGMA) with different distance metrics including Euclidean distance, Pearson’s correlation, and cosine distance. (**C**) K number specification for the optimal discretization levels of pseudo-bulk expression used in Arboretum analysis per 17 neuronal groups. We tested 3-5 discretization levels for each neuronal groups and specified the optimal value based on the observation of Bayesian Information Criteria (BIC, columns E-G) penalized scores and Akaike information criterion (AIC, columns H-J) penalized scores. Red fonts are the final decision.

## METHODS

### Stem Cell Maintenance

Experiments were conducted using the HUES3::Hb9-GFP line (Di Giorgio et al. 2008) (Harvard Stem Cell Institute) or H9 (WA09, WiCell) human embryonic stem cell (hESC) lines in xeno-free, feeder-free conditions. hESCs were maintained at 37°C in 5% CO_2_ in Essential 8 (E8) medium (Lippmann et al., 2014) on Matrigel (WiCell) coated 6-well tissue culture-treated plates and were passaged when 70-80% confluent. Briefly, cells were washed once with phosphate buffered saline (PBS, Invitrogen), then incubated at 37°C in Versene (Invitrogen) for 6 mins. Versene was aspirated, the cells were gently dissociated from the well with fresh E8, and re-plated at a 1:12 seeding ratio. Media was replenished daily (Lippmann et al. 2014).

### Neuromesodermal Progenitor Differentiation

To initiate NMP differentiation, hESCs were washed once with PBS, incubated at 37°C in Accutase (Invitrogen) for 5 mins, singularized by gentle trituration, and quenched with 1 volume E8 media. Following centrifugation for 5 mins at 300g, hESCs were replated onto 35 mm Matrigel-coated plates at a density of 1.5 x 10^5^ cells/cm^2^ in E8 medium with 10 μM ROCK inhibitor (Y27632, Tocris). The media was replaced with Essential 6 (E6) medium (Lippmann et al., 2014) on the following day (Day 0), and then changed to E6 supplemented with 200 ng/mL FGF8b (Peprotech) 24 hours later (Day 1). On Day 2, Hox propagation was initiated by activation of Wnt signaling using NMP media comprised of E6 medium supplemented with 200 ng/mL FGF8b and 3 μM CHIR99021 (Tocris). This constitutes the “Hox timepoint” of 0hrs. At various timepoints, NMPs were collected or differentiated to pCNSPs for scRNAseq experiments and at 120hrs for D/V optimization experiments. For NMPs collected within 24 hours, NMP media was applied directly. Else, cells were sub-cultured at a 2:3 ratio. Briefly, cells were washed once with PBS, incubated in Accutase for 1.5-2 mins, and removed from the surface by gentle pipetting. After centrifugation, cells were gently resuspended in NMP Medium containing 10 μM Y27632 and seeded on 35 mm Matrigel-coated plates. NMP Media was replenished on Day 4. For NMPs collected between 72-96 hours, NMP media was changed directly on Day 5, else cells were sub-cultured again at a 2:3 ratio. The media was replenished daily on Days 7-10 with NMP media containing 30 ng/mL GDF11 (Peprotech) and 1 μM dorsomorphin (Tocris) to stimulate caudal NMP development, with sub-culture on Day 9 at a 1:1 ratio.

### pCNSP Differentiation

To initiate pCNSP differentiation, H9-derived NMPs were cultured for 1 day in spinal progenitor media, comprising of E6 media supplemented with 1 µM retinoic acid (RA; Sigma), 10 µM SB-431542 (Abcam), and 100 nM µL LDN-193189 (Stemgent). Cells were singularized and replated at 5 x 10^5^ cells/cm^2^ in pCNSP media containing 10 μM Y27632 for an additional 2 days. Media was replenished daily.

### D/V Differentiation

pCNSPs were exposed to morphogens for 4 days to initiate D/V patterning. Dorsal progenitors were cultured in E6 media containing 100 μM RA, 1 μM Cyclopamine (Cyc), and BMP4 (Peprotech) at different concentrations and durations. Ventral progenitors were cultured in E6 media containing 100 nM RA, 10 µM SB-431542, 100 nM LDN-193189, and Purmorphamine (Pur; Tocris) and Smoothened Agonists (SAG; Calbiochem) at different concentrations. “High” RA conditions were cultured in 1 μM RA instead of 100 nM RA. Progenitors underwent neuronal differentiation for immunocytochemistry and qPCR studies by switching to maturation media for 5-7 days. Maturation media consisted of E6 containing 1x N2 Supplement (Thermofisher), 50x B27 Supplement (Thermofisher), 1 µM cAMP (Sigma), 10 ng/mL GDNF, 10 ng/mL BDNF, 10 ng/mL NT-3 (Peprotech), and 10 µM DAPT (Tocris). As appropriate, 10 ng/mL BMP7 (Peprotech) was added to maturation media for additional dorsalization. Media was replenished daily.

### Differentiation for Pre-Optimized scRNAseq

Using the HUES3-Hb9-GFP human embryonic stem cell (hESC) line, which fluorescently reports Hb9 (MNX1)^+^ MNs (Di Giorgio et al., 2008), we differentiated NMPs from six timepoints corresponding to 24hrs (H24), 48hrs (H48), 72hrs (H72), 120hrs (H120), 168hrs (H168), or 216hrs (H216) of Hox patterning. Cultures were then switched to E6 media containing 1 μM RA, 2 μM SAG, and 2 μM purmorphamine for 3 days. Progenitors were sub-cultured at a 1:3 ratio and gently resuspended in E6 medium supplemented with 1 μM RA, 100 nM SAG, 100 nM purmorphamine, and 10 μM Y27632 and plated on 35mm Matrigel-coated well-plates for an additional 3 days. Then, the media was switched to E6 medium supplemented with 1 μM RA, 100 nM SAG, 100 nM purmorphamine, and 5 μM DAPT for an additional 5 days, then cryopreserved. Media was replenished daily.

### Differentiation for Optimized scRNAseq

NMPs from six timepoints corresponding to 24hrs (H24), 48hrs (H48), 72hrs (H72), 120hrs (H120), 168hrs (H168), or 216hrs (H216) of Hox patterning were differentiated to pCNSPs. Dorsal progenitors were generated by culturing in E6 media containing 100 μM RA, 1 μM Cyc, 20 ng/mL BMP4 for 1 day, then E6 media containing 100 μM RA and 1 μM Cyc for 3 additional days. Ventral progenitors were generated by culturing in E6 media containing 100 nM RA, 10 µM SB-431542, 100 nM LDN-193189, 0.5 μM Pur, and 0.5 μM SAG for 4 days. For H216R conditions, the RA concentration was increased to 1 μM for both dorsal and ventral differentiations. Progenitors were cryopreserved to synchronize cultures. For neuronal differentiation and maturation prior to sequencing, cells were thawed in maturation media containing 10 μM Y27632 at 5 x 10^5^ cells/cm^2^ overnight, with daily media changes for 6 days. Media was switched to Neurobasal media (Gibco) containing 1x N2 Supplement, 50x B27 Supplement, 1x Glutamax (Thermofisher), 1x PenStrep (Invitrogen), 1 ng/mL laminin (Thermofisher), 1 µM cAMP, 10 ng/mL GDNF, 10 ng/mL BDNF, and 10 ng/mL NT-3 for 14 days. 2 days prior to sequencing, the media was supplemented with 10 µM AraC (Sigma) to eliminate proliferating cells.

### Cryopreservation

To create cryopreserved cell banks for further differentiation or single cell RNA-seq analysis, cells were dissociated in Accutase at 37°C for 30 mins on an orbital shaker, quenched with one volume E6 medium, centrifuged, and gently resuspended in 10% DMSO in E6 medium with 10 μM Y27632. The cells were aliquoted at 1 mL/cryovial and cryopreserved with a CryoMed controlled rate freezer (Thermofisher) using a stepwise cooling program: rapid cooling from room temperature to 4°C, 1°C/min until reaching −60°C, and 10°C/min until reaching −100°C. Cryovials were transferred to a liquid nitrogen dewer for long-term storage.

### Quantitative Real-Time Polymerase Chain Reaction

Total RNA was isolated using TRIzol Reagent (Invitrogen) and cDNA was synthesized using the SuperScript IV First-Strand Synthesis system (Invitrogen) according to the manufacturers’ instructions. TaqMan Gene Expression Assays (**Table S4**) and TaqMan Gene Expression Master Mix (Applied Biosystems) were used on a BioRad CFX96 thermocycler with the following protocol: 50°C for 2 mins; 95°C for 10 mins; 40 cycles of 95°C for 15 seconds and 60°C for 1 minute. Primers used are listed in the STAR Methods Table. Target genes were normalized to RPS18 expression and relative gene expression was calculated using the comparative *ΔCt* method. Fold differences in relative mRNA expression levels of target genes are reported for each gene with standard deviations (*n*=6 biological replicates for each group). Statistical analysis was conducted using JMP-Pro 13 software. Significance was determined using a one-way analysis of variance (ANOVA) with Tukey-Kramer HSD post-hoc for multiple comparisons with a 95% confidence threshold.

### Immunocytochemistry

Cells were fixed in 4% paraformaldehyde for 10 mins, washed thrice in PBS, and blocked in tris-buffered saline (TBS) containing 0.3% triton-x and 5% normal donkey serum (TBSDT) for at least one hour. The cells were incubated in primary antibodies (**Table S4**) diluted in TBSDT overnight at 4°C. After three 15-minute washes in TBS containing 0.3% triton-x, the cells were incubated with Alexa Fluor secondary antibodies (Invitrogen) at a 1:500 dilution in TBSDT for one hour at room temperature. Cells were washed twice in TBS for 15 mins each, counterstained with 300 nM DAPI for 10 mins, and washed once more in TBS prior to mounting with Prolong Gold Antifade Reagent (Life Technologies) as necessary. Images were acquired using a Nikon A1R confocal microscope with Nikon NIS-Elements software and analyzed with NIS-Elements and ImageJ.

### Single Cell Dissociation of Neurons

Cells were singularized for scRNAseq by dissociation with Papain (Worthington). Briefly, cells were washed once with PBS and incubated in papain at 37°C for 1hr on an orbital shaker, then triturated vigorously with a wide-bore pipette. The cell suspension was centrifuged at 300g for 5 minutes, then quenched with ovomucoid solution for 10 minutes at room temperature. Quenched cells were centrifuged, gently resuspended in PBS containing 0.2% BSA and 10 μM Y27632, then passed through a 40 μm cell strainer (Mitenyi Biotec) to remove debris. Cells were quantified and diluted to 700 cells/μL for sequencing.

### Single Cell RNA-Sequencing

Directly after thaw or singularization, ∼3000-5000 cells were targeted for capture from each sample. Transcriptomic profiling was performed using the Chromium Single Cell Gene Expression system (10X Genomics), according to the manufacturer’s recommendations using the Single Cell 30 Reagent v2/v3 kits (10X Genomics). Post-GEM-RT and post-cDNA amplification cleanup were performed using Dynabeads MyOne silane beads (Thermofisher) and SPRIselect (Beckman Coulter) kits respectively. Successful library preparation was confirmed using an Agilent Bioanalyzer (High Sensitivity DNA kit) and Qubit Fluorometer (High Sensitivity dsDNA kit). Experiment data was demultiplexed using the Cell Ranger Single Cell Software Suite, mkfastq command wrapped around Illumina’s bcl2fastq. The MiSeq balancing run was quality controlled using calculations based on UMI-tools (Smith et al., 2017). Samples libraries were balanced for the number of estimated reads per cell and run on an Illumina HiSeq 2500 or NovaSeq 6000 system. Cell Ranger software was then used to perform demultiplexing, alignment, filtering, barcode counting, UMI counting, and gene expression estimation for each sample according to the 10x Genomics documentation (https://support.10xgenomics.com/single-cell-gene-expression/software/pipelines/latest/what-is-cell-ranger). The gene expression estimates from each sample were then aggregated using Cellranger (cellranger aggr) and processed through our data pre-processing pipeline to obtain filtered and normalized expression data.

### Data Pre-Processing

For each of the 6 samples from direct differentiation (GSE###) and 14 samples from multiple generation (GSE###) from our R/C time-series experiment, we filtered out genes that appeared in fewer than 5 cells and cells with fewer than 5000 UMIs from the dataset. Each cell’s expression value was depth normalized to a depth of 5000 followed by variance stabilizing normalization as implemented in the pagoda2 package (Barkas, Nikolas, 2021). We merged the gene expression matrices from each sample into a single matrix while taking the union of the genes from each matrix. The combined matrix is [12,543 cells x 20,598 genes] for the direct differentiation dataset and [49,959 cells x 23,941 genes] for the multiple generation dataset. We transformed the values by taking their square root and standardizing each cell’s expression profile by dividing by the mean expression of a gene in each cell.

### Clustering of Single Cells by HOX Gene Profile

We obtained the expression values of 33 HOX gene paralogs from our normalized matrix to define the *HOX* profile of each cell (**Fig. S4D-G, Table S5**). We clustered the cells based on their *HOX* profile using two different clustering algorithms. Our first approach binarized the *HOX* profiles based on non-zero expression of a HOX gene in a cell and applied k-means clustering with *k* in {5,6,7,8,9,10,11,12,13,14,15,16,17,18,19,20} on these binary profiles. Our second approach applied Louvain clustering on knn graphs of cells with edges weighted by the Euclidean distances between the binary *HOX* expression profiles of each cell. We used 4 different values for the number of neighbors (*n*) in the knn graph, *n* in {20, 30, 40, 50} and searched the resolution parameter of Louvain clustering which controls the number of clusters to identify 5-20 clusters. We started with a resolution of 1 for each desired k. Let k’ be the number of clusters obtained at a resolution of r’. If k’>k, we decreased the resolution by (0.5)*^i^*, else increase the resolution by (0.5)*^i^* where *i* is the iteration of the search. We repeated this until the desired *k* was reached. For each k (the number of clusters), we had 4 different clusterings (for the 4 values of n), and we selected the optimal n based on the lowest Pearson’s correlation between the cluster means. Our rationale was that the Pearson’s correlation would be lowest for the most distinct clusters (**Fig. S4D**). We computed silhouette index for each Louvain clustering and compared this with the kmeans clusters. We used the Louvain clusters for the following analysis as it performed better in the evaluation based on the silhouette index (SI, **Fig. S4E**). To determine k, we examined the patterns of *HOX* genes in each cluster in addition to the SI. Based on SI, k=13 or 14 was optimal. We next annotated each cluster based on the pattern of expression of HOX genes, e.g., a cluster was annotated with a pattern “*HOXA4*+/*HOXA5*-” if *HOXA4* was expressed while *HOXA5* was not. We finally determined the number of clusters to be 13, as it had among the highest SI and captured the most distinct annotation patterns capturing most of the known *HOX* colinear expression patterns (**Fig. S4F-G**). Cluster IDs are rearranged manually by the order of rostrocaudal axis based on the observation of compositions of 9 *HOXA* genes-expressing cells within each cluster.

### Identification Of Primary Clusters

We applied Non-negative Matrix Factorization (NMF) implemented with Alternating Non-negative constrained Least Squares and the active set method (Kim and Park, 2008) with imposing sparsity for the gene-side factors for the identification of primary clusters. This implementation of NMF was shown to have a faster convergence and be more computationally efficient compared to the multiplicative update algorithm originally developed for NMF (Lee and Seung, 1999). Furthermore, based on our comparisons of this algorithm to the Ordinary Least squares implementation in MATLAB, this produced more stable solutions (**Fig. S6A,B**). NMF decomposes an input matrix *X ϵ R^mXn^* into two lower dimensional factors, U and V as 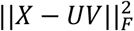 where *U* ∈ *R^mXn^* and *V* ∈ *R^kXn^*. We performed NMF on our merged normalized [cells x genes] matrix with the number of factors/lower dimensions, *k*, to be {5, 10, 15, 20, 25, 30, 35}. We performed k-means clustering on the *U* matrix with k to be the same as the number of the factors for improved clusters. As NMF results in different solutions that are dependent on the starting seed, we applied NMF with 20 different random initializations and assessed the stability based on Jaccard Index of the cluster assignments. Overall, the NMF factorizations were stable (JI ranging from 0.64-0.99) and for each k we took the most stable initialization based on the maximum average Jaccard Indices (JI) between each initialization and the remaining ones (**Fig. S6B**). We used two metrics to determine the number of clusters. First, we extracted 23 well known neural marker genes (**Table S1C**) from our data matrix and calculated the silhouette index (SI) of each clustering solution (**Fig. S6C**). Second, we tested the significance of the difference in expression profiles for each pair of clusters. Briefly, for each k, we first obtained the pseudo-bulk expression of each cluster by taking the mean expression value of a gene across all cells in a cluster. Next, we asked if this expression vector of one cluster was significantly different from another cluster (two-sided paired T-test p<0.05) and counted the proportion of pairs that were significant (**Fig. S6D**). We determined k=25 to be optimal based on SI and the proportion of pairs that were significantly different. To assign cell type identities, we used sample composition of each cluster (**Fig. S6E**) and the relationship between these clusters and the clusters defined using the HOX expression (**Fig. S6F**) to help determine the hindbrain/spinal cord identity. Cluster IDs are rearranged manually by the order of times base on the observation the sample composition.

### Subpopulation Clustering Analysis

We regrouped our 25 primary cell clusters into 17 subgroups based on similarity of the cell types assigned to each cluster: all MNs, somatic MN, visceral MN, Floor Plate (FP)-cranial MN, hindbrain-spinal cord (HB SC) sensory Excitatory, HBSC sensory inhibitory, HB sensory Excitatory, HB sensory inhibitory, SC sensory, HBSC proprioceptive, dl1, dl2, dlA3/dl3, V2a/V3, V2b, RA-dorsal neurons, and RA-ventral neurons. Each subpopulation had between 1,084-11,965 cells (**Table S2**). For each of these group of clusters we aimed to identify robust, high confidence, and fine-grained cell subpopulations indicative of a specific cell type. To this end, we developed a novel clustering pipeline consisting of three steps: (1) ensemble of clusterings, (2) consensus graph generation, and (3) consensus clustering (**Fig. S7A**). We used the V2a/V3 and V2b groups to optimize our pipeline and applied the steps to the remaining 15.

For the first step, we generated a number of clustering solutions to be used for consensus clustering. We compared two different types of clustering approaches for generating the ensemble of clusterings. First, we applied NMF (with the ANLS algorithm) followed by k-means clustering on *U* factors matrix. Second, we applied Louvain clustering with the knn graph estimated using two approaches: (1) a knn graph from pairwise Euclidean distance estimated from a NMF-reduced space of 50 dimensions, and (2) a knn graph estimated using fuzzy simplicial set, used in UMAP and scanpy (Wolf et al., 2018). NMF was applied with the number of factors, k, to be 3-10, each with 10 different random initializations, resulting in 80 different clustering solutions. For both Louvain clustering approaches, we obtained knn graphs with k = {10, 20, 30, 40, 50}, each with 8 different resolutions, {0.01, 0.05, 0.1, 0.3, 0.5, 0.7, 1.0, 1.5}, which in total resulted in 80 different clusterings.

In the second step, we created a consensus graph of cell co-clustering relationship. For every pair of cells, we counted the proportion of times the two cells were in the same cluster (across any of our three clustering approaches, k and resolution), and generated a weighed graph of cells with weights corresponding to this proportion. We generated three types of consensus graphs, one based on NMF only clusterings, one based on Louvain only clusterings and one combining both NMF and Louvain.

In the final step, our goal was to estimate robust cell clusters by clustering the consensus graph. We considered two clustering approaches, one based on NMF and another based on Louvain clustering. For NMF, we considered the number factors in the range of 3-15 and defined cell clusters *l* based on the factor with the largest value for the cell, *i.e. l* = argmax(*U_j_*_1_, *U_j_*_2_, …, *U_jl_*), where *U* ∈ *R^mxl^*, 1 ≤ *j* ≤ *m*. We repeated this procedure 10 times and picked the initialization with the highest Jaccard coefficient with the other clustering solutions for the same k. For Louvain clustering, we extracted the knn graph from the full weighted consensus graph matrix with {10, 20, 30, 40, 50} nearest neighbors and applied clustering at 5 different resolutions as {0.1, 0.3, 0.5, 0.7, 1.0}. We used a metric Delta Consensus Counts (DCC) for measuring the quality of the clusters on the graph. DCC is defined as 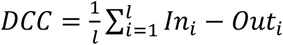 for *l* clusters, where *ln_i_* is the average of edge weights within a cluster *i* and *Out_i_* is the average of edge weights between nodes of cluster *i* to nodes not in cluster *i*. Based on DCC values, we found NMF to be optimal across all three steps, and we applied the same procedure to all other subpopulations (**Fig. S7B**).

Having determined NMF to be the optimal algorithm for our consensus clustering approach, we generated NMF-based consensus clusters for each subpopulation. For all but MNs, we considered k for NMF in step 1, to range from 3-10, with 10 different random initializations, resulting in 80 different clusterings. For the MNs, we used a higher range of k (3-30) because MNs are known to be more complex than others, resulting in 280 clustering solutions. After the consensus graph was generated, we needed to do another round of optimization to select the optimal k on the consensus k. We used a combination of quantitative and qualitative methods. For the quantitative methodology, we used the summation of 3 different evaluation metrics, SI, DCC and stability score (average JI for each pair of clusterings) and shortlisted the top 3-5 best results, which we examined using our qualitative method (**Table S6A**). Here, we manually inspected the block-diagonalness of the clustered consensus graph matrix to avoid over- or under-clustering (**Fig. S7A (iii)**). Based on this procedure, our 17 cluster groups are subdivided into 4-9 fine cell subclusters (**Table S2**). The main paper presents the results of 9 of these groups, the remaining are available in our online resource (https://roy-lab.github.io/subcluster_analysis/). The regional specificities of subclusters were addressed by the observations of sample compositions and *HOX* cluster compositions (**Fig S7C**).

### Comparative Similarity Analysis With Previous Human And Mouse In Vivo Studies

We compared our scRNAseq dataset to two previous *in vivo* studies from human (Rayon et al) and mouse (Delile et al) cells. The raw data from the human and mouse single cell expression studies were downloaded from the Gene Expression Omnibus (GSE171890) and ArrayExpress (E-MTAB-7320) respectively. Each dataset was pre-processed using the same procedure described above and finally merged into a single matrix resulting in 23,179 genes by 47,089 cells for the human dataset and 17,335 genes by 27,725 cells in the mouse dataset. Comparative analysis was restricted to only *SNAP25*^+^ neuronal cells in all datasets, which resulted in 6,026 cells in the mouse dataset and 8,050 cells in the human dataset and 44,487 cells in our dataset. For each dataset, we first grouped cells into cell types based on the expression of marker genes in the Rayon et al. knowledge matrix. We compared our 25 primary clusters (**Fig. 5D**), 17 subpopulations (**Fig. S5A**), and 11 cell types (**Fig. S5B**) defined using the knowledge matrix from Rayon et al. to cell types defined in the mouse (Delile et al) and human (Rayon et al) datasets. For all comparisons, we used 77 marker genes in the knowledge matrix provided by Rayon et al. (using 55 mouse orthologs for Delile et al. obtained from MGI (Bult et al., 2019)) and transcription factors (1,463 genes for human-human and 1,775 genes for human-mouse comparisons) defined by PANTHER and GO. For each type of cell grouping (NMF, subtype or cell types), we obtained a pseudo-bulk expression profile of all marker genes using the mean expression across cells within a group. The similarity between any pair of cell groupings was estimated using the Pearson’s correlation of each group’s pseudo-bulk profiles. A pair of cell groupings were considered matched if there was a high correlation between each row group to one or a few column groups. The best concordance was obtained using the cell type definition of cell groups.

### Identification Of Differentially Expressed Genes (DEG)

*To* define differentially expressed genes (DEGs), we used the intersection of three tests. First, we defined the DEGs for each cluster of each subpopulation as the gene is expressed in more than 50% of the cells in a cell group, and the 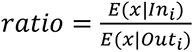 is more than 1.25, where *E*(*x*^*C*) is defined as the number of cells expressing gene *x* in a group of cells *C*. We tested the significance of overlap between all cells expressing each gene and the subcluster by hyper-geometric test. We also applied the Welch’s t-test and Mann-Whitney rank test (Wilcoxon rank-sum test) for differential expression between each cluster and the complement of the cluster of each gene by using the python package diffxpy of the scanpy suite (Wolf et al., 2018). Finally, for the most reliable DEG list, we kept only the DEGs which were significant in all 3 tests (p < 0.05) (**Table S3**).

### Arboretum-Based Identification of Subcluster-Specific Genes

We adapted a previously developed multi-task clustering framework Arboretum (Roy et al., 2013) to find gene modules with similar expression patterns across several subclusters. Arboretum is used to jointly cluster multiple hierarchically related gene expression datasets such that cluster assignments for more similar datasets are more similar. Such relationships could be obtained from phylogenies. The Arboretum framework is based on a generative probabilistic process and has two components: the emission model generates the observed expression measurements at the tips of the tree and is formulated as a mixture of k Gaussians, where k is the number of clusters, and the clusters are related via transition probabilities which model the probabilistic propagation of module assignments from the root of the tree to the tips. In our application of Arboretum to scRNAseq datasets, we first generated pseudo bulk profiles for each cell subcluster, used these to define hierarchies (described next), and finally applied Arboretum to this data with varying values of k. Although the original application of Arboretum models multi-dimensional expression matrices, we used 1-dimensional pseudo-bulk representation of each cell subcluster for efficiency.

To obtain the relationship structure of the cell subclusters, we performed hierarchical clustering based on pairwise distances between pseudo-bulk vectors. For each of the 17 subpopulations, we considered unweighted average distance (UPGMA) with different distance metrics including Euclidean distance, Pearson’s correlation, and cosine distance. We picked the best structure based on the cophenetic correlation coefficient. Different groups were best described by trees from different distance functions (**Table S6B**).

We tested k as {3,4,5} in the Arboretum clustering of each group. The best k was determined using the optimal value across three select metrics: penalized log-likelihood scores, Bayesian information criterion (BIC) penalized score, and Akaike information criterion (AIC) penalized score (**Table S6C**). After clustering, each gene is assigned a cluster assignment in each cell subcluster, which is represented by a vector of discretized expression values across subclusters.

To identify gene sets with combinatorial patterns of expression across the subclusters, we obtained genes that change their cluster assignment across subclusters and applied our previously developed tool, sc*FindTransitioning* (https://github.com/Roy-lab/clade-specific_gene_sets/tree/scFindTransitioning) which makes use of hierarchical clustering. The scFindTransitoning takes a parameter for determining the cluster height to cut the dendrogram. For this analysis, we used the height of 0.05, which was selected based on our previous experience with this tool on other data set. Each of these transitioning gene sets are available at [https://roy-lab.github.io/subcluster_analysis/]. We interpreted these gene sets based on their expression trends as well as known annotations from PANTHER (Thomas et al., 2003) and GO databases (Ashburner et al., 2000).

